# Perturbed structural dynamics underlie inhibition and altered specificity of the multidrug efflux pump AcrB

**DOI:** 10.1101/2020.04.27.063511

**Authors:** Eamonn Reading, Zainab Ahdash, Chiara Fais, Vito Ricci, Xuan Wang Kan, Elizabeth Grimsey, Jack Stone, Giuliano Malloci, Andy M. Lau, Heather Findlay, Albert Konijnenberg, Paula J. Booth, Paolo Ruggerone, Attilio V. Vargiu, Laura J. V. Piddock, Argyris Politis

## Abstract

Resistance-nodulation-division (RND) efflux pumps play a key role in inherent and evolved multidrug-resistance (MDR) in bacteria. AcrB is the prototypical member of the RND family and acts to recognise and export a wide range of chemically distinct molecules out of bacteria, conferring resistance to a variety of antibiotics. Although high resolution structures exist for AcrB, its conformational fluctuations and their putative role in function are largely unknown, preventing a complete mechanistic understanding of efflux and inhibition. Here, we determine these structural dynamics in the presence of AcrB substrates using hydrogen/deuterium exchange mass spectrometry, complemented by molecular modelling, drug binding and bacterial susceptibility studies. We show that the well-studied efflux pump inhibitor phenylalanine-arginine-β-naphthylamide (PAβN) potentiates antibiotic activity by restraining drug-binding pocket dynamics, rather than preventing antibiotic binding. We also reveal that a drug-binding pocket substitution discovered within an MDR clinical isolate, AcrB^G288D^, modifies the plasticity of the transport pathway, which could explain its altered substrate specificity. Our results provide molecular insight into drug export and inhibition of a major MDR-conferring efflux pump and the important directive role of its dynamics.

## Main

AcrB is a homotrimeric integral membrane protein that forms part of the tripartite AcrAB-TolC efflux pump^1^ (Fig. 1a). Energised by the proton-motive force, AcrB transports a broad variety of toxic substances, including antibiotics, outside of the cell through a channel formed by the periplasmic adaptor protein, AcrA, and outer membrane channel, TolC^2,3^. It is constitutively expressed in many pathogenic Gram-negative bacteria and, with its homologues forming the most clinically relevant pumps, has become a target for drug discovery to tackle MDR^4-7^.

**Figure 1.**
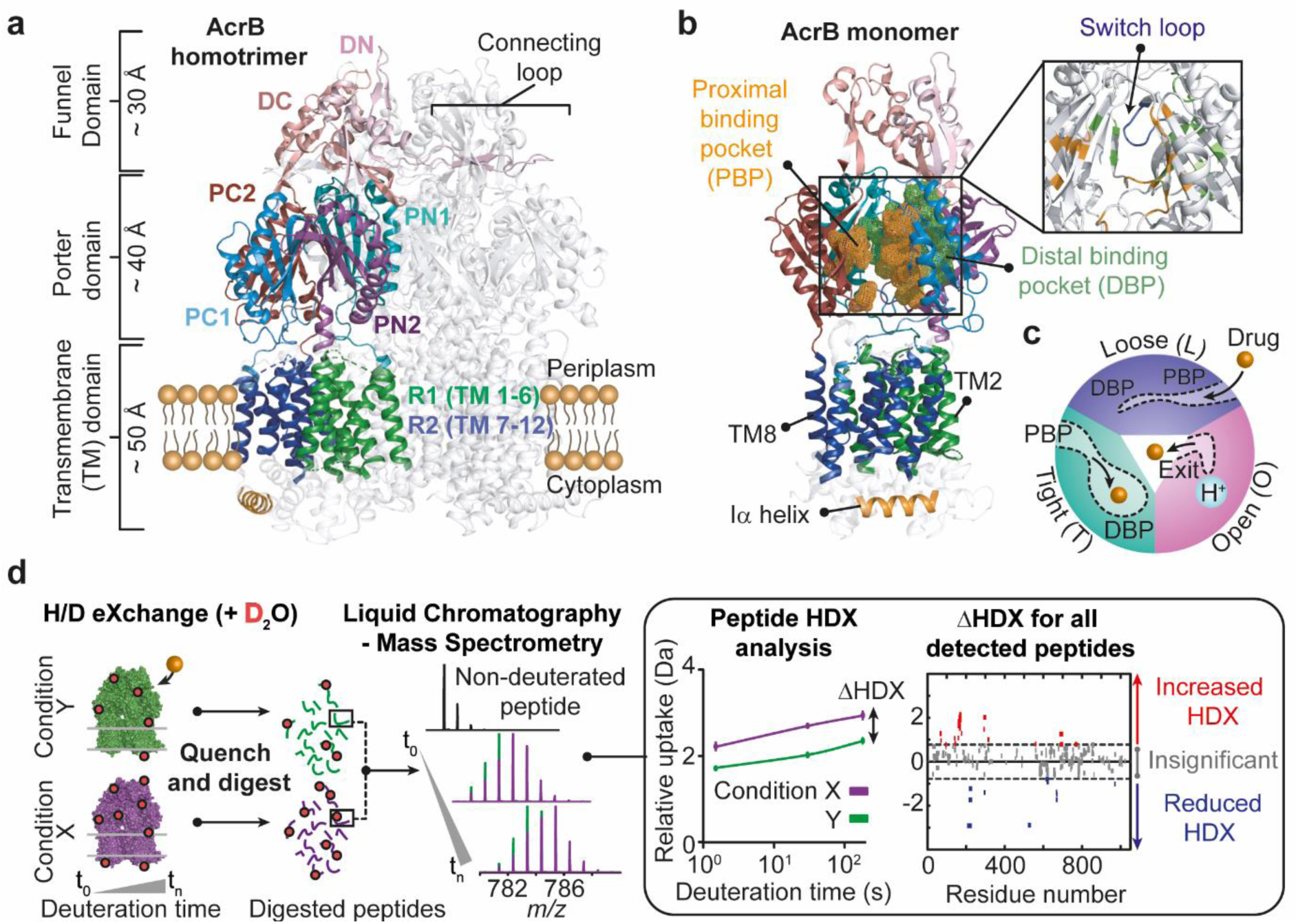
Summary of structure, function, and HDX-MS of AcrB. (**a**) Structure and subdomains of AcrB (PDB:2HRT). (**b**) Highlighted regions of functional interest on AcrB monomeric unit. (**c**) Functional rotation drug efflux mechanism between the three distinct monomer conformations within trimeric AcrB. (**d**) HDX-MS workflow.

Previous structural and biochemical work has enabled drug-binding pockets and efflux pathways for AcrB to be proposed^1,2,7-17^. Drug transport is purported to occur via cooperative rotation between three distinct monomer conformations: loose (L), tight (T) and open (O) (Fig. 1a-c, Supplementary Fig. 1, and Supplementary Table 1). Where, in the L-state, drugs gain access to the proximal binding pocket (PBP) through entrance channels^1,8,18^. Upon a conformational change to the T-state, the drug is then moved towards the distal binding pocket (DBP) before being transported through the exit channel of the periplasmic domain, following a second conformational change from the T- to O-state^19,20^. A switch-loop (^615^FGFAGR^620^) acts to separate the PBP from the DBP; the conformational flexibility of this region is proposed to be essential for the regulation of substrate binding and export (Fig. 1b)^21,22^.

Each AcrB monomer contains a connecting-loop, which protrudes into the Funnel Domain (DC and DN) of the adjacent monomer (Fig. 1a) and has been shown to be important for stabilising the trimer during rotation^23^. During rotation, the transmembrane (TM) region – which consists of the R1 (TM1-6) and R2 (TM7-12) domains connected by the Iα-helix (^520^FEKSTHHYTDSVGGIL^535^) (Fig. 1a-b) - is also postulated to move considerably^14,24^.

Despite the availability of high-resolution structures of AcrB, it has remained challenging to dissect the molecular mechanisms that regulate the impact of naturally occurring MDR mutations, efflux pump inhibition and their synergy with antibiotics. One fundamental aspect of its structure that remains unresolved concerns the role of its structural dynamics, which are often crucial for protein function^25-28^. Here, we use Hydrogen/Deuterium eXchange Mass Spectrometry (HDX-MS) to directly measure changes in AcrB structural dynamics owing to binding of the Ciprofloxacin (CIP) antibiotic and the well-studied PAβN efflux pump inhibitor (EPI)^5,7,15,29^. Investigating both wildtype AcrB (AcrB^WT^) and a recently discovered G288D mutation (AcrB^G288D^), uncovered in a post-therapy MDR clinical isolate of *Salmonella* Typhimurium^30^, we were able to further understand the structural and functional consequences of substrate and inhibitor binding and clinically-relevant mutation.

Our findings reveal that the PAβN EPI restricts the intrinsic motions of the drug-binding pockets as part of its mechanism of action and is effective against both AcrB^WT^ and AcrB^G288D^. We also demonstrate that an EPI can dually bind to AcrB alongside an antibiotic, without affecting its inhibitory action. We discover that an MDR mutation in *acrB* impacts upon the structural dynamics of the efflux translocation pathway, likely contributing to its modified substrate specificity. Structural dynamics therefore have a critical role in the inhibition and substrate specificity of AcrB. Understanding the effects of these dynamics on structure and function is critical for successful assessment of efflux pump mechanism and inhibition, especially in relation to MDR-conferring mutations.

## Results

### Hydrogen/Deuterium eXchange Mass Spectrometry of AcrB

HDX-MS is a solution-based method which can provide molecular level information on local protein structure and dynamics^31-33^. HDX occurs when backbone amides are made accessible to D_2_O solvent through structure unfolding and H-bond breakage; HDX is fast within unfolded regions and slow within stably folded regions (i.e. α-helices, β-sheet interiors), where transient local unfolding events are required for HDX to occur. In order to decipher the impact of drug binding and mutation on AcrB structural dynamics, we performed differential HDX (ΔHDX) analysis between two conditions (e.g. drug-bound and drug-free), which is a sensitive approach for detecting associated structural perturbations between two different protein states (Fig. 1d)^33-35^.

We optimized HDX-MS conditions on *Escherichia coli* (*E. coli*) AcrB solubilized within *n*-Dodecyl-β-D-Maltopyranoside (DDM) detergent micelles achieving 72% peptide coverage (Supplementary Fig. 2). A relative fractional deuterium uptake analysis of AcrB revealed that many of its residues form part of stable structures, inferred from the longest labelling time points (0.5-1 hour) required for substantial deuterium incorporation^32^ (Supplementary Fig. 3). Its most structurally dynamic regions being discovered within the subdomains of the Porter Domain (PC1, PC2, PN1, and PN2) which, notably, host the main drug-binding pockets (Fig. 1a-b). Lack of extensive HDX was observed within the TM domains, likely afforded by their protection within the hydrophobic environment of the detergent micelle.

**Figure 2.**
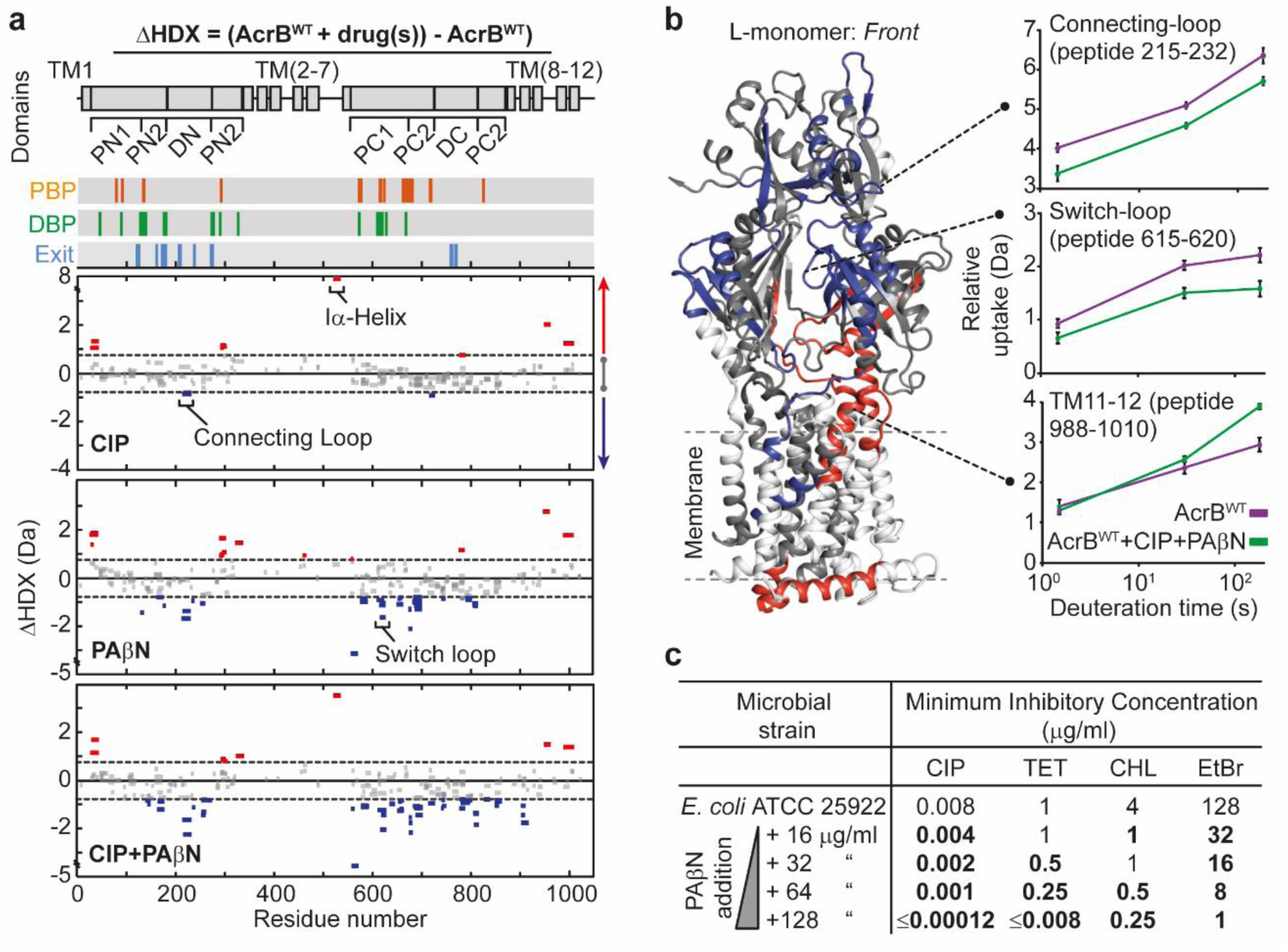
Impact of EPI PAβN on AcrB structural dynamics and function. (**a**) Sum differential HDX (ΔHDX) plots for different drug conditions (ΔHDX = (AcrB^WT^+drug(s)) – AcrB^WT^) for all time points collected. Red signifies peptides with increased HDX (backbone H-bond destabilisation) in drug-bound state and blue represents peptides with decreased HDX (backbone H-bond stabilisation). 98% confidence intervals are shown as grey dotted lines and grey data are peptides with insignificant ΔHDX. All measurements were performed at least in triplicate. All HDX-MS peptide data can be found in Supporting Data Table 2. (**b**) ΔHDX extent for (AcrB^WT^+CIP+PAβN) – AcrB^WT^ is coloured onto the L-state monomer of AcrB (PDB:2HRT) using Deuteros^34^. Connecting-loop from adjacent monomer is included. (**c**) MIC assays of *Escherichia coli* in the presence of inhibitor and antibiotics. Values that demonstrate >2-fold reduction in MIC are in bold. Ciprofloxacin = CIP, Tetracycline = TET, Chloramphenicol = CHL, ethidium bromide = EtBr, and phenylalanine-arginine-β-naphthylamide = PAβN. MIC for PAβN alone is 256 μg/ml.

**Figure 3.**
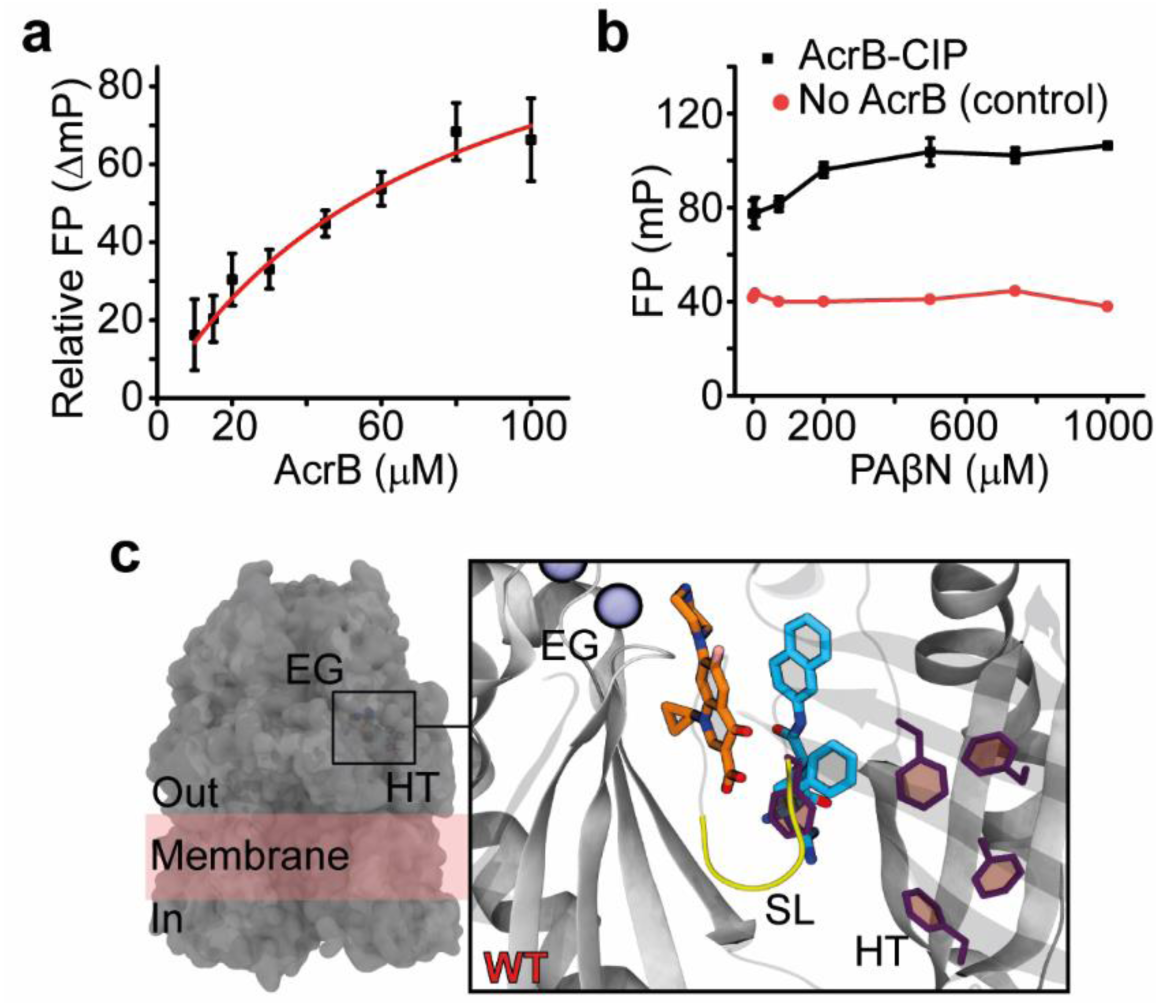
Dual binding of Ciprofloxacin antibiotic and PAβN inhibitor to the DBP of AcrB. (**a**) Binding of CIP by AcrB in the presence of 150 μM of PAβN as determined by a fluorescence polarization assay performed by Su *et al*.^38^. CIP was maintained at 1.5 µM throughout and its emission wavelength measured at 415 nm. Reported data are the average and standard deviation from repeated measurements (n = 3) and were fitted to a hyperbola function (FP = (B_max_*[protein])/(K_D_ + [protein]), R^2^ = 0.97). (**b**) Binding competition assay between PAβN and CIP. PAβN was non-fluorescent in the experimental conditions. Data are average and standard deviation from repeated measurements (n = 3). (**c**) Molecular docking and multi-copy μs-long MD simulations reveal stable interactions of CIP (orange) and PAβN (cyan) to AcrB^WT^ T-state monomer and show their likely binding locations. EG = exit channel gate (blue spheres), SL = switch-loop (yellow), and HT = hydrophobic trap (purple). All computational data can be found in Supplementary Table 2 and Supplementary Fig. 6-8.

### EPI restricts drug-binding pocket dynamics

First, we examined the impact of CIP and PAβN binding on AcrB^WT^. CIP is a licensed antibiotic in clinical use and has been demonstrated to bind to the DBP, PC1/PC2 cleft and central cavity (Supplementary Fig. 1) by MD simulations and X-ray crystallography^15,29^. While PAβN, an EPI and substrate of AcrB, has been shown to bind to similar areas of AcrB as many antibiotics, this binding may be at distinct sites^5,15,29^.

In the presence of CIP only a few regions within the PN2 subdomain, central cavity, R2 domain (TM 7-12) and Iα-helix demonstrated significant differential HDX (Fig. 2a-b), suggesting that antibiotic binding only subtly alters the structural motions of AcrB^WT^. The presence of PAβN gave comparably increased HDX within the PN2 subdomain. However, in stark contrast to CIP, inhibitor binding led to HDX reduction throughout extensive parts of the PC1/PC2 cleft of the drug-binding pockets and within the connecting-loop (Fig. 2a-b). This could signify inhibitor-induced structural stabilisation of the drug-binding pocket entrances.

HDX of the Iα-helix did not significantly change upon addition of PAβN, whereas HDX was substantially increased upon CIP binding (Fig. 2a), possibly because the inhibitor weakens the coupling between the R1 and R2 TM domains. Notably, switch-loop spanning peptides with reduced HDX were detected when PAβN was present, which may reflect a binding interaction and/or structural stabilisation (Fig. 2a-b).

Together, our HDX data support a mode of action of efflux inhibition by which an EPI primarily acts to impart concerted restraint on AcrB structural dynamics, notably restricting the drug-binding pockets, connecting- and switch-loops.

Multi-copy 1-μs long MD simulations (see Methods) of AcrB^WT^ bound to PAβN confirmed that – in agreement with previous literature^6^ – the inhibitor could stably bind to the DBP, straddling the switch-loop and establishing strong interactions with the hydrophobic trap (HT); a peculiar region of the DBP rich in phenylalanines and involved in EPI binding^4,36^ (Supplementary Fig. 4; see also Supplementary Discussion).

**Figure 4.**
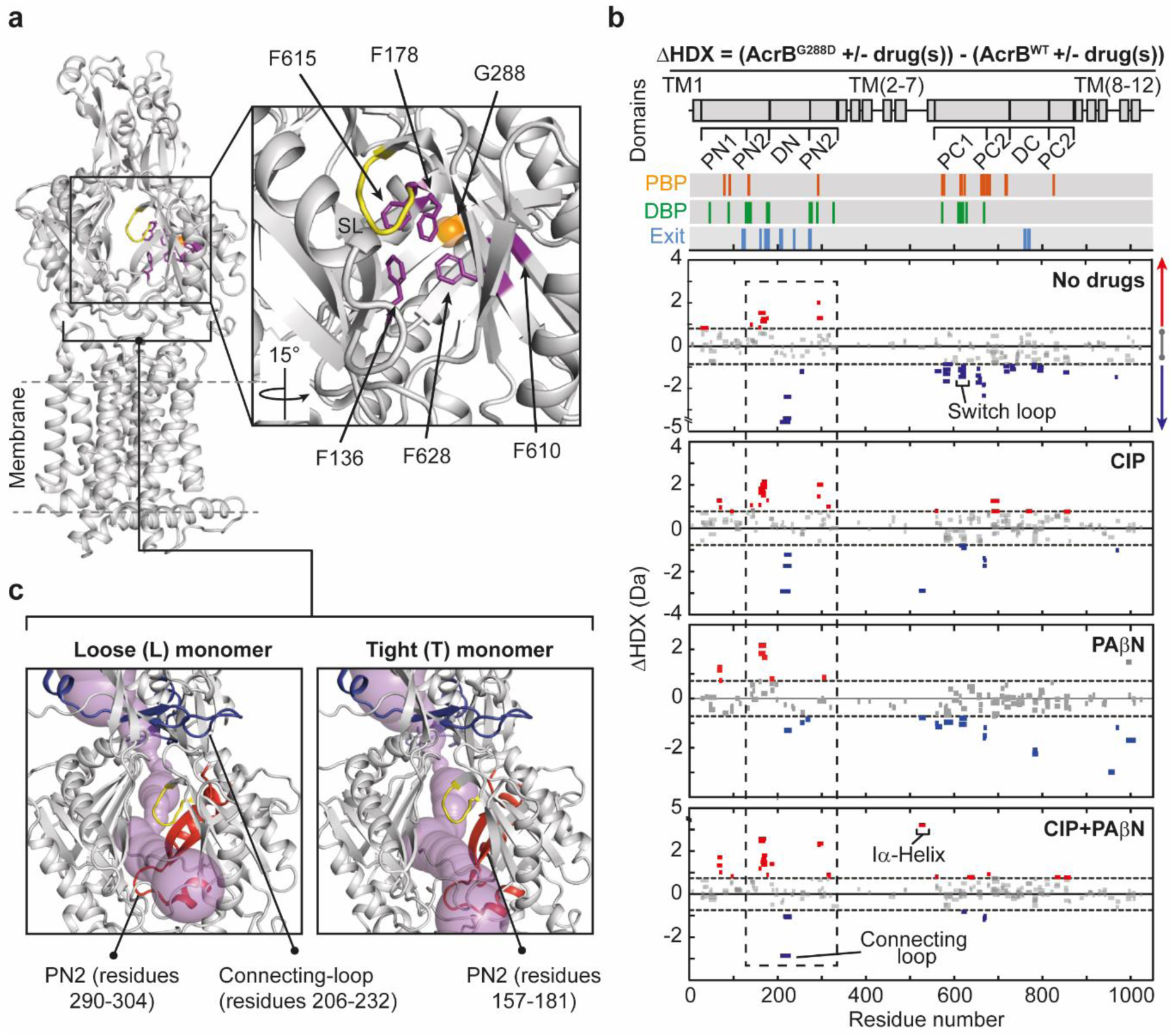
Influence of MDR mutation G288D on AcrB structural dynamics. (**a**) G288 position (orange sphere) amongst switch-loop (SL, yellow) and hydrophobic trap (purple) within the DBP. (**b**) Differential HDX (ΔHDX) plots for AcrB^G288D^ and AcrB^WT^ with and without drugs. All data reported as in Fig. 2. All HDX-MS peptide data can be found in Supporting Data Table 2. (**c**) Shared regions whose HDX are increased (red) and decreased (blue) in a substrate-independent manner (highlighted by a dashed box in Fig. 4b) are shown on AcrB monomers (PDB:2HRT). Connecting-loop from the adjacent monomer is included. Substrate efflux pathway (magenta) for periplasmic entrance was drawn by CAVER^63^.

Comparison between the RMSF profiles of AcrB^WT^-PAβN (T monomer) and apo AcrB^WT^ (L monomer, see Methods) revealed that binding of PAβN is accompanied by an overall rigidification of the protein (Supplementary Fig. 5), which involves large patches of the DBP, PBP, switch-loop, as well as the exit channel gate (EG), CH1, and CH2 channels. In particular: i) residues belonging/adjacent to the switch-loop that directly interact with the inhibitor become more rigid in the presence of PAβN; ii) while the loop itself features moderately enhanced hydration, the nearby segments (612 to 614 and 621 to 624) are overall dehydrated with respect to apo AcrB^WT^. The structural stabilisation occurring upon PAβN binding might prevent local, as well as distal, functional movements that are key to substrate efflux along the transport pathway (Supplementary Fig. 4-5). Thus, we expect the interaction between PAβN and the switch-loop region to be a key factor mediating the mode of action of this EPI.

**Figure 5.**
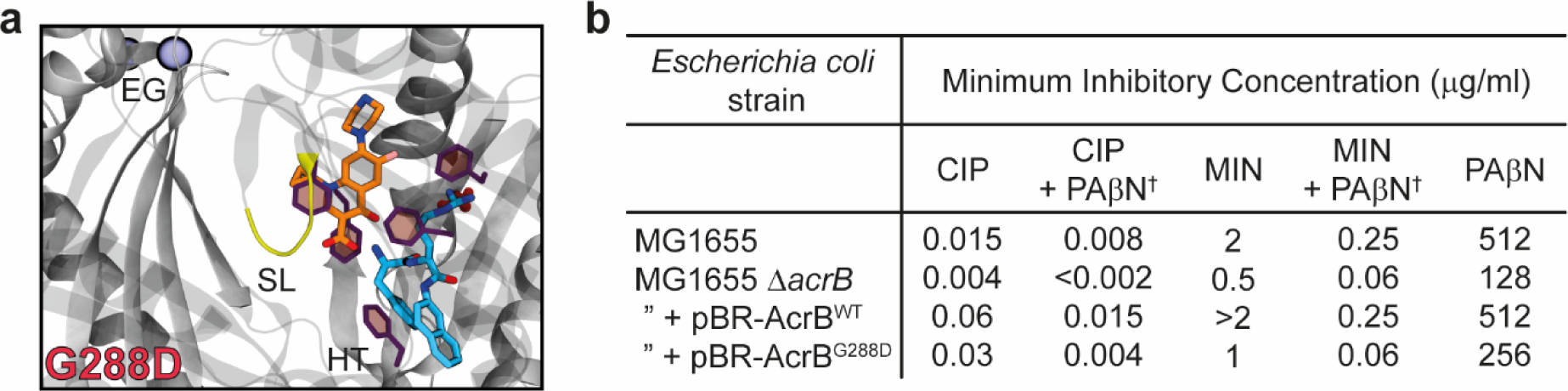
AcrB^G288D^ is inhibited by the EPI PAβN. (**a**) Molecular docking and multi-copy μs-long MD simulations reveal stable interactions of CIP (orange) and PAβN (cyan) to AcrB^G288D^ T-state monomer and show their likely binding locations. The pose and its orientation are the same as shown for AcrB^WT^ in Fig. 3c. EG = exit channel gate (blue spheres), SL = switch-loop (yellow), and HT = hydrophobic trap (purple). All computational data, including binding free energies be found in Supplementary Table 3 and Supplementary Fig. 6, 12-13. (**b**) MIC assays of *Escherichia coli* containing AcrB^WT^ or AcrB^G288D^ in the presence of inhibitors and antibiotics. AcrB was overexpressed in MG1655 *ΔacrB* from a pBR322 plasmid containing its corresponding *acrAB* genes, natural promoter and ‘marbox’ sequence. Minocycline = MIN, Ciprofloxacin = CIP, and phenylalanine-arginine-β-naphthylamide = PAβN. ^†^PAβN was added at a concentration of 50 μg/ml.

Overall, our data are in agreement with a model for inhibitor action, which has been proposed to work by trapping AcrB in a conformation, possibly a T-like state, which prevents adequate functional rotation and substrate transport^16^.

### EPI and antibiotic can dually bind to AcrB

To fully understand EPI action, it is essential to consider its activity in the presence of antibiotic substrates, especially in light of the emerging importance of drug combination therapies for treating bacterial infection^37^. Antibiotic susceptibility assays against *E. coli* confirmed the ability of PAβN to potentiate antibiotic activity (Fig. 2c). PAβN increased antibiotic susceptibility for a range of antimicrobial substrates (CIP, tetracycline (TET), chloramphenicol (CHL), and ethidium bromide (EtBr)) in a substrate-dependent manner, with better effectiveness observed at lower PAβN concentrations for CIP and EtBr than for TET and CHL.

To explore the impact of an antibiotic on efflux inhibition we performed HDX-MS in the presence of both CIP and PAβN (AcrB^WT^-CIP-PAβN). We anticipated that the presence of equimolar CIP may interfere with PAβN binding, thereby impacting its ability to prevent functionality of the transporter through dynamic restrain. This was not the case. The presence of CIP did not alter the action of the PAβN inhibitor, as revealed by the strikingly similar differential HDX profiles for both AcrB^WT^-PAβN and AcrB^WT^-CIP-PAβN (Fig. 2a-b). Consequently, we investigated the possibility that PAβN acts by outcompeting CIP binding to AcrB. To test this, we exploited the innate fluorescence of CIP to perform fluorescence polarization binding and competition assays^38^. Interestingly, we found that CIP binds with similar affinity to both AcrB^WT^ (K_D_ of 74.1 ± 2.6 µM from Su^38^) and a preformed AcrB^WT^-PAβN complex (K_D_ of 76.2 ± 17.4 µM) (Fig. 3a), and that PAβN could not effectively outcompete CIP binding from a preformed AcrB^WT^-CIP complex (Fig. 3b). These data suggest that antibiotic and inhibitor may be able to simultaneously bind at different (sub)sites within the voluminous DBP.

To further support this hypothesis, we performed blind docking and MD simulations on AcrB^WT^-PAβN-CIP (Supplementary Table 2 and Supplementary Fig. 6; see Methods for details). Importantly, both drugs could stably bind to the DBP within the T-state monomer, with PAβN partly occupying the HT and CIP lying in proximity of the PBP/DBP interface (Fig. 3c). Several interactions contribute to stabilize this configuration (Supplementary Fig. 7; see also Supplementary Discussion). The simultaneous binding of CIP and PAβN has similar effects as the binding of the inhibitor only on the flexibility and hydration of the protein (Supplementary Fig. 7-8), in corroboration with our HDX results (Fig. 2a-b).

Overall, our data support the hypothesis that PAβN does not compete or prevent antibiotic binding (competitive inhibition). Instead, we propose that it inhibits efflux by enforcing a more restrained state upon AcrB, thus, reducing the frequency and magnitude of the conformational changes within the substrate translocation path (its effectiveness being substrate dependent).

### MDR-conferring substitution, G288D, impacts drug-binding pocket dynamics

We next turned our attention to the substitution mutation, G288D (AcrB^G288D^), which was found to cause resistance to some drugs (e.g. CIP) in *Salmonella*, but susceptibility to others (e.g. minocycline (MIN))^30^. G288 is a highly conserved residue in Enterobacteriaceae^30^, suggesting an important structural role, and is present aside the HT within the DBP (Fig. 4a). We found that the G288D substitution does not alter the average secondary structure content or thermal stability of AcrB, as judged by circular dichroism (Supplementary Fig. 2). Yet, our HDX analysis revealed that the G288D mutation has a noticeable impact on the structural dynamics of AcrB. We found that, when no drugs were present, the G288D mutation caused increased HDX for several peptides spanning the PN2 region of the protein, but decreased HDX within the PC1/PC2 regions and connecting-loop (Fig. 4b). This suggests that this single-point mutation can cause a long-range change to the structural dynamics of AcrB, possibly reflecting global conformational changes in its substrate-free state.

Markedly, for all three conditions tested (CIP, PAβN, and CIP-PAβN), the G288D substitution consistently caused increased HDX within the PN2 region and decreased HDX of the connecting-loop (Fig. 4b). Signifying that these effects are retained even upon substrate binding and, due to their close structural proximity, may relate to concerted changes to the dynamics of the substrate translocation pathway (Fig. 4c). Whereas, upon substrate binding, HDX within the PC1/PC2 and DC domains of AcrB^G288D^ and AcrB^WT^ became closer in parity (Fig. 4b and Supplementary Figure 9). Although differences were observed; AcrB^G288D^-PAβN had reduced HDX for regions within PC1 and R2 (TM 7-12) domains, in comparison to AcrB^WT^-PAβN.

### PAβN EPI inhibits both wildtype and G288D AcrB

MD simulations of AcrB^G288D^ in the presence of PAβN were performed to better understand how the G288D mutation effects EPI interactions with the drug-binding pockets. PAβN was found to bind to the HT of AcrB^G288D^, directly interacting with the mutated D288 residue through the formation of hydrogen bonds (Supplementary Fig. 10; see also Supplementary Discussion). Interactions with the aromatic residues of the HT involve hydrophobic stacking as well as cation-π attraction, not observed in AcrB^WT^-PAβN (Supplementary Fig. 4) and possibly promoted by the direct interaction of the inhibitor with D288. Similar interactions are also formed with residues of the switch-loop or the surrounding region, in analogy to AcrB^WT^-PAβN. In accordance with HDX-MS, the switch-loop and the surrounding region undergo further dehydration in AcrB^G288D^-PAβN (Supplementary Fig. 11). These data, together with the direct interactions detected between the inhibitor and the region of the switch-loop, support the hypothesis that stabilisation of the latter has a role for the mode of action of PAβN both in AcrB^WT^ and in AcrB^G288D^.

Similar conclusions emerged from the comparison between AcrB^G288D^-CIP-PAβN and AcrB^WT^-CIP-PAβN. Indeed, MD simulations of the former complex revealed that, even upon G288D substitution, CIP and PAβN can stably occupy the DBP at the same time (Fig. 5a and Supplementary Fig. 12-13). Consistent with HDX-MS data (Fig. 4b), further rigidification and dehydration of the switch-loop occur with respect to AcrB^WT^-CIP-PAβN, together with an increase in the hydration and the flexibility of the surrounding region (Supplementary Fig. 12-13; see also Supplementary Discussion). These data advocate that AcrB^G288D^ is inhibited by PAβN in a similar manner as AcrB^WT^.

The above findings were supported by bacterial susceptibility assays on *E. coli* containing overexpressed AcrB^G288D^; in the presence of PAβN inhibitor there was increased antibiotic susceptibility to MIN and CIP antibiotics for both AcrB^WT^ and AcrB^G288D^ (Fig. 5b). AcrB^G288D^ was more susceptible to PAβN than AcrB^WT^. The decreased susceptibility of AcrB^G288D^ to CIP, discovered within *Salmonella* clinical isolates previously^30^, was not recapitulated in our assays using the laboratory *E. coli* strain MG1655. However, the associated increased susceptibility to MIN was observed (Fig. 5b), supporting that G288D has a profound impact on AcrB substrate specificity within both *E. coli* and *Salmonella*.

## Discussion

In summary, we found that binding of an EPI, PAβN, restricts AcrB dynamics and could not be outcompeted by an antibiotic whose activity it potentiates. Endorsing the hypothesis that RND-pump inhibitors act through an “altered-dynamics” mechanism, which despite interfering little with substrate binding, impacts significantly on the kinetics of the functional cycle.

Furthermore, we reveal that a clinically relevant drug-pocket substitution, G288D^30^, impacts the structural dynamics of AcrB - with the substrate translocation pathway being modified in a substrate-independent manner. Our previous MD simulations^30^ on AcrB^G288D^ demonstrate that the G288D substitution increased the gyration radius and the hydration at and around the DBP. This could subsequently lead to the observed allosteric action on farther AcrB regions, particularly in the perturbation of the substrate efflux pathway (Fig. 4c). In turn, these changes may alter the energetic barrier for substrate binding and transport during functional rotation and, consequently, be the cause for its altered substrate specificity.

More generally, we anticipate that these findings will be important not only for defining how mutant and substrate interactions impact AcrB structural dynamics, which are hard to elucidate from biochemical and high-resolution structural data alone, but also for understanding their general role in modulating multidrug binding and efflux.

## Acknowledgements

Work at King’s College London was supported by a UKRI Future Leaders Fellowship (MR/S015426/1) to E. R., the Wellcome Trust (109854/Z/15/Z) to A.P., a Wellcome Trust Investigator Award 214259 to P.J.B., and a King’s Health Partners R&D Challenge Fund through the MRC Confidence in Concept Grant (MC_PC_15031) to A.P., E.R., and P.J.B.. Work at the University of Cagliari received support from the National Institutes of Allergy and Infectious Diseases project number AI136799 (C.F., G.M., P.R., and A.V.V.) and the Innovative Medicines Initiatives Joint Undertaking under grant agreement number 115525 (C.F., G.M., P.R., and A.V.V.), resources that are composed of financial contribution from the European Union 7th Framework Programme (FP7/2007-2013) and EFPIA companies in kind contribution. Work at the University of Birmingham by L.J.V.P., X.W.K., J.S. and V.R. was supported by MRC grant, (MR/P022596/1) and E.M.G. by an MRC iCASE studentship (MR/N017846/1). We thank Bram Snijders, Peter Faull and Shahid Mehmood (Francis Crick Institute, London) for the use of their Thermo Scientific Q Exactive UHMR hybrid Quadrupole-orbitrap mass spectrometer.

## Author Contributions

E.R., Z.A., A.V.V, L.J.V.P, and A.P. designed the research. E.R., Z.A., A.M.L, and H.F. performed all experiments and analyses, except for molecular modelling and bacterial susceptibility assays. C.F., G.M., and A.V.V. carried out docking, molecular dynamics and performed related analyses. X.W.K., V.R., J.S., and E.M.G. performed bacterial susceptibility assays and analysis. A.K. and Z.A. performed UHMR Native MS measurements and analysis. E.R., P.J.B., P.R., A.V.V, L.J.V.P, and A.P. supervised the project. E.R., Z.A., and A.P. wrote the manuscript with input from the other authors.

## Competing interests

The authors declare no competing financial interests.

## Methods

### Plasmid construction

An overexpression plasmid containing AcrB with a C-terminal 6xHistidine tag (AcrB-6xHis) was constructed from a pET15b-AcrB-sGFP plasmid from Reading *et al*.^39^. Briefly, the sGFP sequence was deleted and a 6xHis-tag placed at the C-terminus of AcrB followed by a stop codon using the Q5^®^ site-directed mutagenesis kit (New England Biolabs) - AcrB contains two Histidine residues at its C-terminus, therefore, this construct resulted in AcrB having an 8xHistidine tag. The G288D mutation was then generated from this pET15b-AcrB-6xHis plasmid using the Q5^®^ site-directed mutagenesis kit (New England Biolabs). All constructs were verified by DNA sequencing (Eurofins MWG).

pBR322-AcrB plasmids were generated for bacterial susceptibility assays. Briefly, pBR322 was linearized with HindIII and EcoRI restriction enzymes (New England Biolabs). *acrAB* genes with its natural promoter, including the ‘marbox’ sequence, was then amplified from K-12 *Escherichia coli* chromosomal DNA (Zyagen Labs) and cloned into the pBR322 vector using In-Fusion^®^ HD cloning (Takara Bio). A 6xHistidine tag sequence was included in the reverse primer to provide a 6xHis tag at the C-terminus of AcrB (pBR-AcrB^WT^). The G288D mutation was generated from this pBR-AcrB^WT^ plasmid using the Q5^®^ site-directed mutagenesis kit (New England Biolabs). All constructs were verified by primer walking DNA sequencing (Eurofins MWG).

### AcrB overexpression and purification

pET15b-AcrB-6xHis plasmid containing AcrB wildtype (AcrB^WT^) or G288D mutant (AcrB^G288D^) was transformed into C43(DE3)Δ*acrB::KanR E. coli* cells. 7 ml of an overnight LB culture was added to 1 L of pre-warmed LB culture containing 100 µg/ml ampicillin and 30 µg/ml kanamycin and grown at 37 °C until an OD of 0.6-0.8 was reached. The culture was induced with 1 mM IPTG and grown for 16-18 hrs at 18 °C. At which point the cells were harvested by centrifugation at 4,200 x *g* for 30 min and washed with ice cold phosphate buffer saline (PBS).

Cell pellets were immediately resuspended in buffer A (50 mM sodium phosphate, pH 7.4, 300 mM sodium chloride) and supplemented with a protease inhibitor tablet (Roche), 100 μM PMSF, 1 µl Benzonase, and 5 mM beta-mercaptoethanol (β-ME). The cell suspension was then passed twice through a microfluidizer processor (Microfluidics) at 25,000 psi and 4 °C. Insoluble material was removed by centrifugation at 20,000 x *g* for 30 min at 4 °C. Membranes were then pelleted from the supernatant by centrifugation at 200,000 x *g* for 1 hr at 4 °C. Membrane pellets were resuspended to 40 mg ml^-1^ in ice-cold buffer A supplemented with a protease inhibitor tablet (Roche) and 100 μM PMSF, and homogenized using a Potter-Elvehjem Teflon pestle and glass tube.

AcrB was extracted from homogenized membranes by overnight incubation with 1 % (w/v) *n*-dodecyl-β-D-maltoside (DDM) detergent (Anatrace) at 4 °C with gentle agitation. Insoluble material was then removed by centrifugation at 100,000 x *g* for 1 hr at 4 °C. The sample was then filtered through a 0.22 μm filter (Fisher Scientific) and loaded onto a 1 ml HiTrap column (GE Healthcare) equilibrated in buffer B (50 mM sodium phosphate, pH 7.4, 300 mM sodium chloride, 20 mM imidazole, 10 % (w/v) glycerol, 0.03% (w/v) DDM). The column was washed with 5 CVs of buffer B and then with 10 CVs of buffer C (50 mM sodium phosphate, pH 7.4, 300 mM sodium chloride, 20 mM imidazole, 10 % (w/v) glycerol, 1% (w/v) octyl glucose neopentyl glycol (OGNG) (Generon)) – an OGNG detergent wash facilitates the removal of bound lipopolysaccharide (LPS), as determined previously^40^. The column was then washed with 20 CVs of Buffer B containing 50 mM imidazole. AcrB was then eluted with Buffer B containing 500 mM imidazole and directly injected onto a Superdex 75 10/600 GL size exclusion chromatography (SEC) column (GE Healthcare) equilibrated in buffer D (50 mM sodium phosphate, pH 7.4, 150 mM sodium chloride, 10 % (w/v) glycerol, 0.03 % (w/v) DDM). Peak fractions eluted from the SEC column containing pure AcrB were pooled and spin filtered before being flash frozen and stored at -80 °C. SDS-PAGE electrophoresis was used to assess AcrB purification and protein concentration was calculated using a Cary 300 Bio UV-Vis spectrophotometer (Varian) with a calculated extinction coefficient^41^ of ε_280_ = 89,730 M^−1^ cm^-1^.

### Circular dichroism spectroscopy

Circular dichroism spectroscopy (CD) spectra was recorded with an Aviv Circular Dichroism spectrophotometer, Model 410 (Biomedical Inc., Lakewood, NJ, USA), with specially adapted sample detection to eliminate scattering artefacts. Multiple CD scans were averaged, the buffer background subtracted, and zeroed and minimally smoothed using CDTool^42^. A final protein concentration of 0.5-1.5 mg ml^-1^ was used in a quartz rectangular or circular Suprasil demountable cell (Hellma Analytics). For thermal protein unfolding the mean residue ellipticity at 222 nm was monitored with increasing temperature.

### Native mass spectrometry

Purified AcrB was buffer exchanged into MS buffer (200 mM ammonium acetate, pH 7.4, 0.03% (w/v) DDM) using a centrifugal buffer exchange device (Micro Bio-Spin 6, Bio-Rad) as previously described^43^. Native mass spectrometry experiments were performed either on a Synapt G2-Si mass spectrometer (Waters) or a Thermo Scientific Q Exactive UHMR hybrid Quadrupole-orbitrap mass spectrometer.

For experiments on the Synapt G2-Si mass spectrometer the instrument settings used were: 1.5 kV capillary voltage, source temperature of 25 °C, argon trap collision gas, 180 V trap collision voltage, 120 V cone voltage, and 50 V source offset. Data was processed and analysed using MassLynx v.4.1 (Waters).

Native mass spectrometry data on a Thermo Scientific Q Exactive UHMR hybrid Quadrupole-orbitrap mass spectrometer was acquired at resolving power 8750 at *m/z* 400 in the *m/z* range 2000 – 30.000. The instrument was optimized for transmission and desolvation of integral membrane proteins. Critical parameters throughout were relative pressure of 6, capillary temperature 250 °C, S-Lens RF level 0, in-source trapping 200 and the HCD energy was 300%. Data were analyzed by use of Xcalibur software and Biopharma Finder 3.1 (both Thermo Fisher Scientific). Deconvoluted spectra were acquired using Bipharma Finder 3.1 in sliding window mode using the following settings: Output mass range 10000-100000 Da, deconvolution mass tolerance 10 ppm, sliding window merge tolerance 30 ppm and minimal number of detected intervals.

### Preparation of ligands for hydrogen/deuterium mass spectrometry

Ciprofloxacin (CIP) antibiotic and Phe-Arg-β-naphthylamide dihydrochloride (PAβN) inhibitor were both purchased from Sigma Aldrich. Stock concentrations of CIP (10 mg/ml) and PAβN (10 mg/ml) were prepared in water. As demonstrated previously^44^, a primary consideration before carrying out HDX-MS is for ligands with known dissociation constants to ensure close to full binding to the target protein under deuterium exchange conditions; particularly for low affinity ligands (dissociation constants, kD, in the μM range or lower). CIP and PAβN both possess moderate affinity binding to AcrB within DDM detergent micelles (CIP with a k_D_ of 74.1 ± 2.6 μM, as measured by fluorescence polarization^38^ and PAβN with a k_D_ of 15.72 ± 3.0 μM, as measured by surface plasmon resonance^45^). To achieve sufficient ligand binding saturation under deuterium exchange conditions AcrB was first incubated in the presence of saturating concentrations of ligand for 30 minutes on ice, before dilution into deuterated buffer containing saturating concentrations of ligand (740 μM, ∼1000:1 ligand to AcrB ratio) for deuterium exchange experiments.

### Hydrogen/deuterium mass spectrometry

Hydrogen/deuterium mass spectrometry (HDX-MS) was performed on an HDX nanoAcquity ultra-performance liquid chromatography (UPLC) Synapt G2-Si mass spectrometer system (Waters Corporation). Optimized peptide identification and peptide coverage for AcrB was performed from undeuterated controls. The optimal sample workflow for HDX-MS of AcrB was as follows: 5 μl of AcrB (15 μM) was diluted into 95 μl of either buffer D or deuterated buffer D at 20 °C. After fixed times of deuterium incubation samples were mixed with a formic acid-DDM quench solution to provide a quenched sample at pH 2.5 and final 0.075 % (w/v) DDM concentration. The quenched sample was then injected onto an online Enzymate™ pepsin digestion column (Waters) in 0.1 % formic acid in water (at a flow rate of 200 μl/min) maintained at 20 °C. The peptic fragments were trapped onto an Acquity BEH C18 1.7 μM VANGUARD pre-column (Waters) for 3 min. The peptic fragments were then eluted using an 8-40% gradient of 0.1 % formic acid in acetonitrile at 40 μl/min into a chilled Acquity UPLC BEH C18 1.7 μM 1.0 × 100 mm column (Waters). The trap and UPLC columns were both maintained at 0°C. The eluted peptides were ionized by electrospray into the Synapt G2-Si mass spectrometer. MS^E^ data was acquired with a 20 to 30 V trap collision energy ramp for high-energy acquisition of product ions. Leucine enkephalin was used for lock mass accuracy correction and the mass spectrometer was calibrated with sodium iodide. The on-line Enzymate™ pepsin column was washed three times with pepsin wash (1.5 M Gu-HCl, 4 % MeOH, 0.8 % formic acid), as recommended by the manufacturer, and a blank run was performed between each sample to prevent significant peptide carry-over between runs.

All deuterium time points and controls were performed in triplicate. Sequence identification was performed from MS^E^ data of digested undeuterated samples of AcrB using the ProteinLynx Global Server 2.5.1 software. The output peptides were then filtered using DynamX (v. 3.0) using the following filtering parameters: minimum intensity of 1000, minimum and maximum peptide sequence length of 4 and 25 respectively, minimum MS/MS products of 1, minimum products per amino acid of 0.12, and a maximum MH^+^ error threshold of 15 ppm. Additionally, all the spectra were visually examined and only those with suitable signal to noise ratios were used for analysis. The amount of relative deuterium uptake for each peptide was determined using DynamX (v. 3.0) and was not corrected for back exchange^32^. The relative fractional uptake (RFU) was calculated from *RFU*_*a*_ *= [Y*_*a,t*_*/(MaxUptake*_*a*_ *x D)]*, where *Y* is the deuterium uptake for peptide *a* at incubation time *t*, and *D* is the percentage of deuterium in the final labelling solution.

Confidence intervals for differential HDX-MS (ΔHDX) measurements of any individual time point were then determined according to Houde *et al*.^46^ using Deuteros software^34^. There was no correlation found between ΔHDX values and their standard deviations (R^2^ = 0.09). Only peptides which satisfied a ΔHDX confidence interval of 98 % were considered significant. All ΔHDX AcrB structure figures were generated from the data using Deuteros in-house source code/software^34^ and Pymol^47^. All data and meta-data are reported in Supporting Data Tables 1-2.

### Bacterial susceptibility assays

Antimicrobial susceptibility testing of all wildtype and mutant strains in the absence or presence of PAβN was determined in triplicate by the broth microdilution (BMD) method and the standardized agar doubling-dilution method as recommended by the European Committee on Antimicrobial Susceptibility Testing (EUCAST). EUCAST guidelines were followed conforming to ISO 20776-1:2006^48,49^. Antibiotics and EPIs were made up and used according to the manufacturer’s instructions. *E. coli* ATCC 25922 was used as the control strain.

### Fluorescence Polarization

AcrB ligand binding was determined using fluorescence polarization (FP) assays as performed by Su *et al*.^38^. An AcrB-PAβN protein complex stock was prepared; to ensure a loaded complex, AcrB and 150 μM PAβN was incubated for 2 hours at 25 °C before titrating with 1.5 μM CIP (k_D_ of PAβN is 15.72 ± 3.0 μM, as measured by surface plasmon resonance^45^). AcrB protein titration experiments were performed in ligand binding solution (50 mM sodium phosphate, 150 mM sodium chloride, 10 % (v/v) glycerol, 1.5 μM CIP, 150 μM PAβN, 0.03 % (w/v) DDM, pH 7.4). FP measurements were taken after incubation for 5 min for each corresponding protein concentration to ensure that the binding has reached equilibrium. Ligand binding data was fit to a hyperbola function (FP = (B_max_*[protein])/(k_D_ + [protein])) as performed previously by Su *et al*.^38^ using ORIGIN Ver. 7.5. (OriginLab Corporation, Northampton, MA, USA).

A FP competition assay was performed by titrating increasing concentrations of PAβN (0-1000 µM) to a AcrB-CIP preformed complex concentration adjudged from the binding data (1.5 µM CIP and 45 µM AcrB) - CIP and AcrB was maintained at the same concentration during all PAβN titrations. FP measurements were taken after incubation for 5 min for each corresponding protein concentration to ensure that the binding has reached equilibrium.

Each data point was an average of 15 FP measurements and each titrations series was performed three times. The absorption spectra of PAβN from 350 to 500 nm exhibited that PAβN absorbs light at 350 and 370 nm. The excitation wavelength of CIP at 415 nm does not excite PAβN, therefore PAβN can be considered as a non-fluorescent ligand within these experiments. DDM detergent concentration was consistent to eliminate possible changes in polarization by drug–DDM micelle interactions.

### Molecular docking

A blind docking campaign was first performed using Autodock Vina^50^. As done in Atzori et al.^51^, a rectangular search space of size 125Å × 125Å × 110Å enclosing the whole portion of the protein potentially exposed to ligands was adopted. The exhaustiveness parameter, related to the extent of the exploration within the search space, was set to 8192 (∼1000 times the default 8) in order to improve the sampling of docking poses within the large box used (∼64 times the default 30Å × 30Å × 30Å). Flexibility of both partners was considered indirectly, by employing multiple conformations in ensemble docking runs^52^. For both CIP and PAβN, 10 representative molecular conformations were obtained from 1 μs-long molecular dynamics simulations of the compounds in presence of explicit solvent^53^ (data available at www.dsf.unica.it/translocation/db). Namely, a cluster analysis of the trajectories of the ligands was performed as described in Malloci et al.^53^, setting the number of cluster representatives to 10.

For the wildtype receptor (AcrB^WT^), 10 X-ray asymmetric high-resolution structures (with PDB IDs: 2GIF, 2DHH, 2J8S, 3W9I, 4DX5, 4DX7, 4U8V, 4U8Y, 4U95, 4U96) were considered, most bearing a substrate bound to the transporter. For the G288D variant of AcrB (AcrB^G288D^), we also employed 10 structures, namely the homology models derived on top of the AcrB^WT^ X-ray structures mentioned above. Regarding the homology modelling protocol, the sequence of the G288D variant was first generated by manually modifying the FASTA file of the corresponding amino acid sequence of *E. coli* AcrB retrieved from the Uniprot database (Uniprot Id: P31224). Next, 100 homology models were generated for each template with the Modeller 9.21^54^ software. The variable target function method was used to perform the optimization, and the best model (that is the one with the highest value of the MOLPDF function) was employed in docking calculations.

The ensemble docking campaign resulted in several hundred poses per ligand, most of which were located inside the distal binding pocket of the monomer in the T state (DBP_T_), which is the putative binding site for the recognition of low molecular mass compounds such as those studied here^11^ (see Supplementary Fig. 1 and Supplementary Table 1). Because most docking poses were concentrated in this region, we performed a second docking campaign using a grid of 30Å × 30Å × 30Å and centered at DBP_T_. Next, we performed a cluster analysis of the docking poses using as a metric the heavy-atoms Root Mean Square Deviation (RMSD) of the substrate (setting the cutoff to 3 Å), which returned respectively 11, 9, 15 and 17 different poses for the AcrB^WT^–PAβN, AcrB^WT^–CIP–PAβN, AcrB^G288D^–PAβN, AcrB^G288D^–CIP–PAβN complexes (Supplementary Tables 2, 3).

### Molecular dynamics simulations

All of the 52 complexes selected from docking runs were subjected to all-atom molecular dynamics (MD) simulations (each of 1 μs in length) performed with the AMBER18 package^55^. Protomer-specific protonation states of AcrB were adopted following previous work^56^: residues E346 and D924 were protonated only in the L and T protomers, while residues D407, D408, and D566 were protonated only in the O protomer, of AcrB. The topology and the initial coordinate files were created using the LEaP module of the AMBER18 package. The proteins were embedded in a mixed bilayer patch composed of 1-palmitoyl-2-oleoyl-*sn*-glycero-3-phosphoethanolamine (POPE) and 1-palmitoyl-2-oleoyl-*sn*-glycero-3-phosphoglycerol (POPG) in a 2/1 ratio, for a total of 660 lipid molecules symmetrically distributed in the two leaflets of the bilayer. The whole system was solvated with a 0.15 M aqueous NaCl solution. The AMBER force field protein.fb15^57^ was used to represent the protein; lipid17 (http://ambermd.org/GetAmber.php) parameters were used for the POPE molecules; the TIP3PFB model was employed for water^58^. The GAFF force-field parameters^59^ for CIP and PAβN were taken from Malloci et al^53^.

Each system was first subjected to a multi-step structural relaxation via a combination of steepest descent and conjugate gradient methods using the *pmemd* program implemented in AMBER18, as described in previous publications^4,29,56^. The systems were then heated from 0 to 310 K in two subsequent MD simulations: i) from 0 to 100 K in 1 ns under constant-volume conditions and with harmonic restraints (k = 1 kcal·mol^-1^·Å^-2^) on the heavy atoms of both the protein and the lipids; ii) from 100 to 310 K in 5 ns under constant pressure (set to a value of 1 atm) and with restraints on the heavy atoms of the protein and on the z coordinates of the phosphorous atoms of the lipids to allow membrane rearrangement during heating. As a final equilibration step, a series of 20 equilibration steps, each of which was 500 ps in duration (total 10 ns), with restraints on the protein coordinates, were performed to equilibrate the box dimensions. These equilibration steps were carried out under isotropic pressure scaling using the Berendsen barostat, whereas a Langevin thermostat (collision frequency of 1 ps^−1^) was used to maintain a constant temperature. Finally, production MD simulations of 1 μs were performed under an isothermal-isobaric ensemble for each system. A time step of 2 fs was used for all runs before production, while the latter runs were carried out with a time step of 4 fs after hydrogen mass repartitioning^60^.

During the MD simulations, the lengths of all the R–H bonds were constrained with the SHAKE algorithm. Coordinates were saved every 100 ps. The Particle mesh Ewald algorithm was used to evaluate long-range electrostatic forces with a non-bonded cut-off of 9 Å.

### Post-processing of MD trajectories

MD trajectories were analyzed using either in-house *tcl* and *bash* scripts or the *cpptraj* tool of AMBER18. Figures were prepared using gnuplot 5.0^61^ and VMD 1.9.3^62^. All the calculations with the exception of the cluster analysis were performed on the conformations taken from the most populated conformational cluster (representing the most sampled conformation of the complex along the production trajectories) along the last 200 ns of the production runs.

#### Cluster analysis

Clustering of the ligand trajectory was carried out using the average-linkage hierarchical agglomerative clustering method implemented in *cpptraj* and employing an RMSD cut-off of 3 Å calculated on all the heavy atoms of the ligand.

#### System Stability

The RMSDs of the protein and of the substrates were calculated using *cpptraj* after structural alignment of each trajectory. Namely, we calculated the Cα-RMSD of the protein with respect to the initial (docking) structure after alignment of the whole trimer. The RMSDs of the substrates were calculated with respect to the corresponding structure of the selected docking pose, as well as with respect to the last frame of the MD trajectory. In particular, to evaluate the magnitude of the displacements and reorientations of the substrates during the simulations, their RMSDs were calculated upon alignment of the T monomer of the protein to the reference frame.

#### System Flexibility

The Root Mean Square Fluctuations (RMSFs) of the protein were calculated using *cpptraj* after structural alignment of each trajectory as described in the previous paragraph.

#### Hydration properties

Residue-wise average numbers of waters within the first (second) hydration layer were calculated with *cpptraj* using a distance cut-off of 3.4 (5.0) Å between the nitrogen of the protein and the water oxygens.

#### Comparison with HDX-MS data

RMSFs and hydration properties of each system were compared with a proper reference state according to the current knowledge about the most likely conformations assumed by AcrB in the absence of ligands or complexed with substrates and inhibitors^16^. For instance, to account for conformational changes of AcrB induced by inhibitor binding, PAβN-bound and apo AcrB structures were considered in their T and L state, respectively. The T state was also considered for systems containing both PAβN and CIP (AcrB^WT^-CIP-PAβN and AcrB^G288D^-CIP-PAβN), hypothesizing their stability in this conformation, as evidenced by the RMSDs analyses conducted on our trajectories (Supplementary data Figs. 4, 5, 8, 9). The list of reference states used for each analysis are reported in Supplementary data Table 4.

#### Stabilizing interactions and hydrogen bonds

Stabilizing interactions were analysed by considering residues within 3.5 Å of each substrate in the last 300 ns of the MD trajectories. Hydrogen bonds were identified through geometrical criteria, using a cut-off of 3.2 Å for the distance between donor and acceptor atoms and a cut-off of 135° for the donor-hydrogen-acceptor angle. Such analyses were conducted through *in-house* tcl scripts.

## Notes

### Competing Interest Statement

The authors have declared no competing interest.

## References

1 Du, D. et al. Multidrug efflux pumps: structure, function and regulation. Nat. Rev. Microbiol. 16, 523–539, doi:10.1038/s41579-018-0048-6 (2018).

2 Du, D. et al. Structure of the AcrAB-TolC multidrug efflux pump. Nature 509, 512–515, doi:10.1038/nature13205 (2014).

3 Symmons, M. F., Bokma, E., Koronakis, E., Hughes, C. & Koronakis, V. The assembled structure of a complete tripartite bacterial multidrug efflux pump. Proc. Natl. Acad. Sci. U.S.A. 106, 7173–7178, doi:10.1073/pnas.0900693106 (2009).

4 Sjuts, H. et al. Molecular basis for inhibition of AcrB multidrug efflux pump by novel and powerful pyranopyridine derivatives. Proc. Natl. Acad. Sci. U.S.A. 113, 3509, doi:10.1073/pnas.1602472113 (2016).

5 Kinana, A. D., Vargiu, A. V., May, T. & Nikaido, H. Aminoacyl β-naphthylamides as substrates and modulators of AcrB multidrug efflux pump. Proc. Natl. Acad. Sci. U.S.A. 113, 1405–1410, doi:10.1073/pnas.1525143113 (2016).

6 Vargiu, A. V., Ruggerone, P., Opperman, T. J., Nguyen, S. T. & Nikaido, H. Molecular Mechanism of MBX2319 Inhibition of Escherichia coli AcrB Multidrug Efflux Pump and Comparison with Other Inhibitors. Antimicrob. Agents Chemother. 58, 6224, doi:10.1128/AAC.03283-14 (2014).

7 Yu, E. W., McDermott, G., Zgurskaya, H. I., Nikaido, H. & Koshland, D. E., Jr. Structural basis of multiple drug-binding capacity of the AcrB multidrug efflux pump. Science 300, 976–980, doi:10.1126/science.1083137 (2003).

8 Zwama, M. & Yamaguchi, A. Molecular mechanisms of AcrB-mediated multidrug export. Res. Microbiol. 169, 372–383, doi:10.1016/j.resmic.2018.05.005 (2018).

9 Murakami, S., Nakashima, R., Yamashita, E., Matsumoto, T. & Yamaguchi, A. Crystal structures of a multidrug transporter reveal a functionally rotating mechanism. Nature 443, 173–179, doi:10.1038/nature05076 (2006).

10 Eicher, T. et al. Transport of drugs by the multidrug transporter AcrB involves an access and a deep binding pocket that are separated by a switch-loop Proc. Natl. Acad. Sci. U.S.A. 109, 5687 (2012).

11 Nakashima, R., Sakurai, K., Yamasaki, S., Nishino, K. & Yamaguchi, A. Structures of the multidrug exporter AcrB reveal a proximal multisite drug-binding pocket. Nature 480, 565–569, doi:10.1038/nature10641 (2011).

12 Seeger, M. A. et al. Structural asymmetry of AcrB trimer suggests a peristaltic pump mechanism. Science 313, 1295–1298, doi:10.1126/science.1131542 (2006).

13 Murakami, S., Nakashima, R., Yamashita, E. & Yamaguchi, A. Crystal structure of bacterial multidrug efflux transporter AcrB. Nature 419, 587–593, doi:10.1038/nature01050 (2002).

14 Eicher, T. et al. Coupling of remote alternating-access transport mechanisms for protons and substrates in the multidrug efflux pump AcrB. Elife 3, doi:10.7554/eLife.03145 (2014).

15 Yu, E. W., Aires, J. R., McDermott, G. & Nikaido, H. A periplasmic drug-binding site of the AcrB multidrug efflux pump: a crystallographic and site-directed mutagenesis study. J. Bacteriol. 187, 6804–6815, doi:10.1128/JB.187.19.6804-6815.2005 (2005).

16 Wang, Z. et al. An allosteric transport mechanism for the AcrAB-TolC multidrug efflux pump. eLife 6, e24905, doi:10.7554/eLife.24905 (2017).

17 Ruggerone, P., Murakami, S., Klaas, M. P. & Attilio, V. V. RND Efflux Pumps: Structural Information Translated into Function and Inhibition Mechanisms. Curr. Top. Med. Chem. 13, 3079–3100, doi:10.2174/15680266113136660220 (2013).

18 Oswald, C., Tam, H.-K. & Pos, K. M. Transport of lipophilic carboxylates is mediated by transmembrane helix 2 in multidrug transporter AcrB. Nat. Commun. 7, 13819, doi:10.1038/ncomms13819 (2016).

19 Matsunaga, Y. et al. Energetics and conformational pathways of functional rotation in the multidrug transporter AcrB. eLife 7, e31715, doi:10.7554/eLife.31715 (2018).

20 Vargiu, A. V. et al. Water-mediated interactions enable smooth substrate transport in a bacterial efflux pump. Biochim. Biophys. Acta, Gen. Subj. 1862, 836–845, doi:10.1016/j.bbagen.2018.01.010 (2018).

21 Muller, R. T. et al. Switch Loop Flexibility Affects Substrate Transport of the AcrB Efflux Pump. J. Mol. Biol. 429, 3863–3874, doi:10.1016/j.jmb.2017.09.018 (2017).

22 Ababou, A. & Koronakis, V. Structures of Gate Loop Variants of the AcrB Drug Efflux Pump Bound by Erythromycin Substrate. PLOS ONE 11, e0159154, doi:10.1371/journal.pone.0159154 (2016).

23 Fang, J., Yu, L., Wu, M. & Wei, Y. Dissecting the function of a protruding loop in AcrB trimerization. J. Biomol. Struct. Dyn. 31, 385–392, doi:10.1080/07391102.2012.703065 (2013).

24 Zwama, M. et al. Hoisting-Loop in Bacterial Multidrug Exporter AcrB Is a Highly Flexible Hinge That Enables the Large Motion of the Subdomains. Front. Microbiol. 8, 2095–2095, doi:10.3389/fmicb.2017.02095 (2017).

25 Karplus, M. & Kuriyan, J. Molecular dynamics and protein function. Proc. Natl. Acad. Sci. U.S.A. 102, 6679, doi:10.1073/pnas.0408930102 (2005).

26 Shaw, D. E. et al. Atomic-level characterization of the structural dynamics of proteins. Science 330, 341–346, doi:10.1126/science.1187409 (2010).

27 Bhabha, G. et al. A Dynamic Knockout Reveals That Conformational Fluctuations Influence the Chemical Step of Enzyme Catalysis. Science 332, 234, doi:10.1126/science.1198542 (2011).

28 Campbell, E. et al. The role of protein dynamics in the evolution of new enzyme function. Nat. Chem. Biol. 12, 944–950, doi:10.1038/nchembio.2175 (2016).

29 Vargiu, A. V. & Nikaido, H. Multidrug binding properties of the AcrB efflux pump characterized by molecular dynamics simulations. Proc. Natl. Acad. Sci. U.S.A. 109, 20637 (2012).

30 Blair, J. M. A. et al. AcrB drug-binding pocket substitution confers clinically relevant resistance and altered substrate specificity. Proc. Natl. Acad. Sci. U.S.A. 112, 3511–3516, doi:10.1073/pnas.1419939112 (2015).

31 Trabjerg, E., Nazari, Z. E. & Rand, K. D. Conformational analysis of complex protein states by hydrogen/deuterium exchange mass spectrometry (HDX-MS): Challenges and emerging solutions. Trends Anal. Chem. 106, 125–138, doi: https://doi.org/10.1016/j.trac.2018.06.008 (2018).

32 Wales, T. E., Eggertson, M. J. & Engen, J. R. Considerations in the analysis of hydrogen exchange mass spectrometry data. Methods Mol. Biol. 1007, 263–288, doi:10.1007/978-1-62703-392-3_11 (2013).

33 Masson, G. R. et al. Recommendations for performing, interpreting and reporting hydrogen deuterium exchange mass spectrometry (HDX-MS) experiments. Nat. Methods 16, 595–602, doi:10.1038/s41592-019-0459-y (2019).

34 Lau, A. M. C., Ahdash, Z., Martens, C. & Politis, A. Deuteros: software for rapid analysis and visualization of data from differential hydrogen deuterium exchange-mass spectrometry. Bioinformatics 35, 3171–3173, doi:10.1093/bioinformatics/btz022 (2019).

35 Chalmers, M. J., Busby, S. A., Pascal, B. D., West, G. M. & Griffin, P. R. Differential hydrogen/deuterium exchange mass spectrometry analysis of protein-ligand interactions. Expert Rev. Proteomics 8, 43–59, doi:10.1586/epr.10.109 (2011).

36 Nakashima, R. et al. Structural basis for the inhibition of bacterial multidrug exporters. Nature 500, 102, doi:10.1038/nature12300 (2013).

37 Tyers, M. & Wright, G. D. Drug combinations: a strategy to extend the life of antibiotics in the 21st century. Nat. Rev. Microbiol. 17, 141–155, doi:10.1038/s41579-018-0141-x (2019).

38 Su, C. C. & Yu, E. W. Ligand-transporter interaction in the AcrB multidrug efflux pump determined by fluorescence polarization assay. FEBS Lett. 581, 4972–4976, doi:10.1016/j.febslet.2007.09.035 (2007).

39 Reading, E. et al. The role of the detergent micelle in preserving the structure of membrane proteins in the gas phase. Angew. Chem. Int. Ed Engl. 54, 4577–4581, doi:10.1002/anie.201411622 (2015).

40 Reading, E. et al. The Effect of Detergent, Temperature, and Lipid on the Oligomeric State of MscL Constructs: Insights from Mass Spectrometry. Chem. Biol. 22, 593–603, doi:10.1016/j.chembiol.2015.04.016 (2015).

41 Gasteiger, E. et al. in The Proteomics Protocols Handbook (ed John M. Walker) 571–607 (Humana Press, 2005).

42 Lees, J. G., Smith, B. R., Wien, F., Miles, A. J. & Wallace, B. A. CDtool-an integrated software package for circular dichroism spectroscopic data processing, analysis, and archiving. Anal. Biochem. 332, 285–289, doi:10.1016/j.ab.2004.06.002 (2004).

43 Laganowsky, A., Reading, E., Hopper, J. T. & Robinson, C. V. Mass spectrometry of intact membrane protein complexes. Nat. Protoc. 8, 639–651, doi:10.1038/nprot.2013.024 (2013).

44 Chandramohan, A. et al. Predicting Allosteric Effects from Orthosteric Binding in Hsp90-Ligand Interactions: Implications for Fragment-Based Drug Design. PLOS Comput. Biol. 12, e1004840, doi:10.1371/journal.pcbi.1004840 (2016).

45 Mowla, R., Wang, Y., Ma, S. & Venter, H. Kinetic analysis of the inhibition of the drug efflux protein AcrB using surface plasmon resonance. Biochim. Biophys. Acta, Biomembr. 1860, 878–886, doi:https://doi.org/10.1016/j.bbamem.2017.08.024 (2018).

46 Houde, D., Berkowitz, S. A. & Engen, J. R. The utility of hydrogen/deuterium exchange mass spectrometry in biopharmaceutical comparability studies. J. Pharm. Sci. 100, 2071–2086, doi:10.1002/jps.22432 (2011).

47 Schrodinger, LLC. The PyMOL Molecular Graphics System, Version 1.3r1 (2010).

48 EUCAST Definitive Document E.DEF 3.1, June 2000: Determination of minimum inhibitory concentrations (MICs) of antibacterial agents by agar dilution. Clin. Microbiol. Infect. 6, 509–515, doi:10.1046/j.1469-0691.2000.00142.x (2000).

49 ISO. Clinical laboratory testing and in vitro diagnostic test systems — Susceptibility testing of infectious agents and evaluation of performance of antimicrobial susceptibility test devices — Part 1: Reference method for testing the in vitro activity of antimicrobial agents against rapidly growing aerobic bacteria involved in infectious diseases. (International Organization for Standardization, Geneva, Switzerland, 2006).

50 Trott, O. & Olson, A. J. AutoDock Vina: improving the speed and accuracy of docking with a new scoring function, efficient optimization, and multithreading. J. Comput. Chem. 31, 455–461, doi:10.1002/jcc.21334 (2010).

51 Atzori, A. et al. Identification and characterization of carbapenem binding sites within the RND-transporter AcrB. Biochim. Biophys. Acta, Biomembr. 1861, 62–74, doi:10.1016/j.bbamem.2018.10.012 (2019).

52 Amaro, R. E. et al. Ensemble Docking in Drug Discovery. Biophys. J. 114, 2271–2278, doi:10.1016/j.bpj.2018.02.038 (2018).

53 Malloci, G. et al. A Database of Force-Field Parameters, Dynamics, and Properties of Antimicrobial Compounds. Molecules 20, doi:10.3390/molecules200813997 (2015).

54 Šali, A. & Blundell, T. L. Comparative Protein Modelling by Satisfaction of Spatial Restraints. J. Mol. Biol. 234, 779–815, doi:https://doi.org/10.1006/jmbi.1993.1626 (1993).

55 D.A. Case et al. AMBER 2018. (2018).

56 Ramaswamy, V. K., Vargiu, A. V., Malloci, G., Dreier, J. & Ruggerone, P. Molecular Rationale behind the Differential Substrate Specificity of Bacterial RND Multi-Drug Transporters. Sci. Rep. 7, 8075, doi:10.1038/s41598-017-08747-8 (2017).

57 Wang, L.-P. et al. Building a More Predictive Protein Force Field: A Systematic and Reproducible Route to AMBER-FB15. J. Phys. Chem. B 121, 4023–4039, doi:10.1021/acs.jpcb.7b02320 (2017).

58 Gonzalez-Salgado, D. & Vega, C. A new intermolecular potential for simulations of methanol: The OPLS/2016 model. J. Chem. Phys. 145, 034508, doi:10.1063/1.4958320 (2016).

59 Wang, J., Wolf, R. M., Caldwell, J. W., Kollman, P. A. & Case, D. A. Development and testing of a general amber force field. J. Comput. Chem. 25, 1157–1174, doi:10.1002/jcc.20035 (2004).

60 Hopkins, C. W., Le Grand, S., Walker, R. C. & Roitberg, A. E. Long-Time-Step Molecular Dynamics through Hydrogen Mass Repartitioning. J. Chem. Theory Comput. 11, 1864–1874, doi:10.1021/ct5010406 (2015).

61 Williams, T. & Kelley, C. gnuplot 5.0. (2017).

62 Humphrey, W., Dalke, A. & Schulten, K. VMD: visual molecular dynamics. J. Mol. Graph 14, 33-38, 27-38 (1996).

63 Jurcik, A. et al. CAVER Analyst 2.0: analysis and visualization of channels and tunnels in protein structures and molecular dynamics trajectories. Bioinformatics 34, 3586–3588, doi:10.1093/bioinformatics/bty386 (2018).

