## Supplementary Information for "Perturbed structural dynamics underlie inhibition and altered specificity of the multidrug efflux pump AcrB"

### Supplementary discussion

In this section we report a more in-depth analysis of our MD simulations, focusing on the details of the interactions established between AcrB and the two ligands CIP and PAβN. For each system, we present the stable binding poses, which in some cases reflect ligand orientations and ligand-AcrB interactions that are representative of more than one MD replica.

**AcrB$^{WT}$-PAβN**

According to our MD simulations, an important contribution to the stabilization of PAβN in AcrB$^{WT}$ comes from the hydrophobic trap (HT), whose residues are involved in stacking with the β-naphthylamide moiety of the inhibitor (see Pose 1 in Supplementary Fig. 4 for a representation of the binding mode). Importantly, these interactions also involve residues of the switch loop (such as F617) or of the adjacent regions. These findings, in agreement with previous literature[5,6], support the hypothesis that the stabilization of the switch-loop could be key to the mode of action of PAβN. This loop is also involved in the formation of hydrogen bonds with the amino group of the compound. Additional hydrogen bonds are formed by its polar groups with polar and acid residues of the DBP, including E130, K131 (involved in interactions with the guanidino group of PAβN) and Q176 (interacting with the carbonyl group).

Important findings on the stabilization of the switch loop come from the comparison of the hydration properties of AcrB$^{WT}$-PAβN and apo AcrB$^{WT}$ (Supplementary Fig. 6). Indeed, despite the relevant difference in the timescales of all-atom MD simulations and typical HDX kinetics (see Methods), and while the switch-loop itself moderately enhanced hydration, the nearby segments (residues 612 to 615 and 620 to 624) are overall dehydrated with respect to the apo form (Supplementary Fig. 6). Considerable dehydration and rigidification are also observed for part of the PBP and the CH2 entrance (Supplementary Fig. 6), consistently with HDX-MS data (Fig. 2a-b and Supplementary Table 1).

In agreement with previous studies[5,6], all of the binding modes found for AcrB$^{WT}$-PAβN (Supplementary Fig. 4) feature the β-naphthylamide moiety of the inhibitor within the HT and interacting with the switch loop or nearby residues. Additional common interactions involve the aromatic rings and the amino group of the inhibitor and the PBP/DBP interface, including residues of the PN1 subdomain (such as S46, S128 and E130). Residues of this region are involved in either stacking interactions with the phenyl ring of PAβN (Supplementary Fig. 4, Pose 1, 2) or hydrogen bonds with the amino group of the inhibitor (Pose 3). Furthermore, residues belonging to segment 130-134 also interact with the guanidino group of PAβN in two representative poses (Pose 1, 3), with additional stabilization provided by Q176 and proximal residues. The other pose (Pose 2) is characterized by a different orientation of the guanidino group of the inhibitor, located in the upper part of the DBP and involved in interactions with D276 and nearby residues.

**AcrB$^{WT}$-CIP-PAβN**

Interactions stabilizing CIP and PAβN in AcrB$^{WT}$ include hydrogen bonds between the two substrates (see Pose 1 in Supplementary Fig. 5 for a representation of the binding pose), as well as between them and the protein (e.g. between the guanidino group of PAβN and residues E130 and D174, or between R620 and the carboxylic and carbonyl group in CIP). Additional stabilization comes from stacking of aromatic rings, formed by PAβN with CIP and F615. Importantly, the direct interaction between the inhibitor and segments proximal to the switch-loop, present in AcrB$^{WT}$-PAβN (Supplementary Fig. 4, Pose 1), is preserved also in the presence of CIP.

The comparison of the hydration properties of AcrB$^{WT}$-CIP-PAβN (Supplementary Fig. 7) and AcrB$^{WT}$-PAβN (Supplementary Fig. 6) reveals an analogous dehydration of the residues of the binding pockets (exception made for some residues involved in interactions with the compounds, such as E173, N174 and F615 in AcrB$^{WT}$-CIP-PAβN - see Supplementary Fig. 5, Pose 1) and similar variations in the region surrounding the switch-loop (Supplementary Fig. 6, 7). Such region, involved in interactions with the substrates in both AcrB$^{WT}$-PAβN (Supplementary Fig. 4, Pose 1) and AcrB$^{WT}$-CIP-PAβN (Supplementary Fig. 5, Pose 1), is indeed considerably rigidified in both systems. This is associated to a dehydration of the segments adjacent to the loop, significantly marked in AcrB$^{WT}$-CIP-PAβN (Supplementary Fig. 7), in agreement with HDX-MS data (Fig. 2a-b).

A common trait of the binding modes found for this ternary complex is the presence of direct interactions between the two substrates, through the formation of hydrogen bonds (involving, in all poses, the carboxylic group of CIP) as wells as stacking of the aromatic rings (Supplementary Fig. 5, Pose 1, 3, 4). In particular, while both CIP and PAβN are located inside the DBP in three representative poses (Pose 1, 3, 4), a different binding mode is possible where CIP is located within the PBP behind the switch loop (Pose 2).

Although some differences are present, comparison of the binding regions reveals several shared traits. Firstly, interactions of at least one substrate with the HT and (the region proximal to) the switch loop are preserved. Typically, such interactions involve π-stacking with the aromatic groups of PAβN, although cation-π interactions were also observed in Pose 3 (involving e.g. F178 and the amino group of PAβN). In Pose 2 and 4, additional stacking interactions are found between CIP and the switch loop and the nearby residues.

With the exception of Pose 3, another conserved trait is related to the interaction with the PBP/DBP interface. Several contacts with residues of this region (such as S46, T87, S128 and adjacent residues) are formed by CIP in Poses 1 and 4, while in Pose 2 PAβN is involved in hydrogen bonds and polar interactions with E130 and nearby residues. In Pose 3, in which interactions with the PBP/DBP interface are not detected, several contacts are formed by CIP with polar and acidic residues present in PN2 portion of the DBP, including E152 and Q151.

**AcrB$^{G288D}$-PAβN**

As in the wild type protein, also in the mutant a significant contribution to the stabilization of PAβN comes from the residues of the HT and of the region around the switch-loop. Such residues are indeed involved in stacking with the aromatic groups of the inhibitor, as well as in cation-π interactions with its guanidino and amino groups (see Pose 1 in Supplementary Fig. 8 for a representation of the binding pose). These interactions, possibly promoted by the hydrogen bond formed by the amino group of PAβN with D288, were not detected in AcrB$^{WT}$-PAβN (Supplementary Fig. 4, Pose 1); thus, they provide an additional contribution to the stabilization of the inhibitor specifically for the G288D mutant. Additional contacts not observed in AcrB$^{WT}$-PAβN are formed with part of the PC1/PC2 cleft (such as L668), while interactions with the PBP/DBP interface, present in AcrB$^{WT}$-PAβN, are not retained.

Although small differences in the flexibility of the binding sites emerged from the comparison of the RMSFs of AcrB$^{G288D}$-PAβN and AcrB$^{WT}$-PAβN (both considered in the T state, see Methods) (Supplementary Fig. 10), higher hydration levels were detected in AcrB$^{G288D}$-PAβN within the DBP and particularly at residues around D288, which include F178 and adjacent residues in PN2. These findings are in good agreement with HDX-MS data (Fig. 4b).

A comparison of the binding poses in AcrB$^{G288D}$-PAβN (Supplementary Fig. 8) reveals a strong contribution to the stabilization of the system from stacking of the aromatic groups of the inhibitor with residues of the HT, in analogy to our findings in AcrB$^{WT}$-PAβN (Supplementary Fig. 4) and AcrB$^{WT}$-CIP-PAβN (Supplementary Fig. 5). Additional stabilization comes from cation-π interactions involving the guanidino group of PAβN and residues belonging or proximal to the switch-loop, as well as the amino group of the inhibitor and F178 in Pose 1. Moreover, contacts are formed between PAβN and the substituted residue D288, involved e.g. (in Pose 1) in hydrogen bonds with the amino group of PAβN.

In both poses, stabilizing interactions further involve residues of the PC1/PC2 cleft, such as π -stacking with aromatic groups of PAβN (Pose 1, 2) or hydrogen bonds between the amino group of the inhibitor and the backbone of residues P669 and A670 (Pose 2).

**AcrB$^{G288D}$-CIP-PAβN**

As in AcrB$^{WT}$, CIP and PAβN are involved in direct interactions through hydrogen bonds that involve the amino group of the inhibitor and the carboxylic group of CIP (see Pose 1 in Supplementary Fig. 9 for a representation of the binding pose). The amino group of the inhibitor is additionally involved in cation-π interactions with residue F178, while its guanidino group is oriented towards D288. Additional stabilization comes from the stacking of the aromatic groups of PAβN with the lower part of the HT (F136, Y327) and the cleft (segment 668-670). CIP is also involved in stacking with residues close to the switch-loop (such as F615), as well as in interactions with hydrophobic residues proximal to the HT (I277, V612). In analogy to AcrB$^{WT}$-CIP-PAβN (Supplementary Fig. 5, Pose 1), therefore, contacts with the switch-loop are retained, but the interaction with the PBP/DBP interface is weakened. Moreover, stabilizing interactions also involve some residues of the PC1/PC2 cleft representing the entering gate towards the PBP, as well as the mutated residue D288.

From the comparison of the flexibility and hydration properties of AcrB$^{G288D}$-CIP-PAβN and AcrB$^{WT}$-CIP-PAβN (Supplementary Fig. 11), it emerges that the switch-loop is considerably more rigid and dehydrated in the mutant. A net increase in hydration and flexibility is also detected for part of the PN2 portion of the DP (including segment 178-182, involved in interactions with the substrates – see Supplementary Fig. 9, Pose 1 and Supplementary Fig. 11). These data agree with the stabilization of the switch-loop and the increase in hydration of PN2 emerged from HDX-MS analyses (Fig. 4b).

As for AcrB$^{WT}$-CIP-PAβN, direct interactions between the two substrates that involve the carboxylic and carbonyl group of CIP are present in both the binding modes detected in

AcrB$^{G288D}$-CIP-PAβN (Supplementary Fig. 9). In addition, π-stacking between one or both substrates and the HT and the switch loop were detected. Further stabilization is provided by cation-π interactions established between the amino group of PAβN and F178 (Pose 1). Importantly, D288 also contributes to stabilize the complex by forming hydrogen bonds with PAβN (Pose 1) or CIP (Pose 2). Another common feature of both poses is the π-stacking formed with residues of the PC1/PC2 cleft. A major difference regards instead the interaction with the PBP/DBP interface, which is indeed present only in Pose 2 and involves the phenylalanine and arginine moieties of PAβN and the carboxylic group of CIP. In Pose 1 PAβN is located inside the HT and CIP interacts with regions proximal to the switch loop and to the upper part of the DBP (including, for example, residues I277 and segment 178-182).

### Supplementary Figures

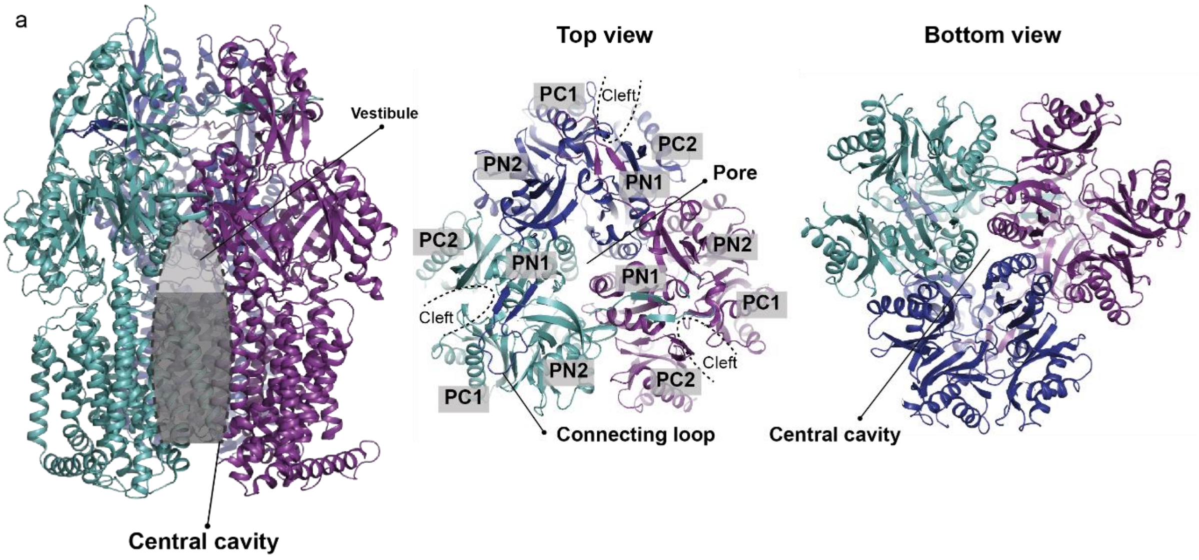


**Supplementary Figure 1. AcrB central cavity and trimer architecture.** The central cavity is shown by a dashed dotted line and a dark grey area, and the vestibule is shown by a light grey area.

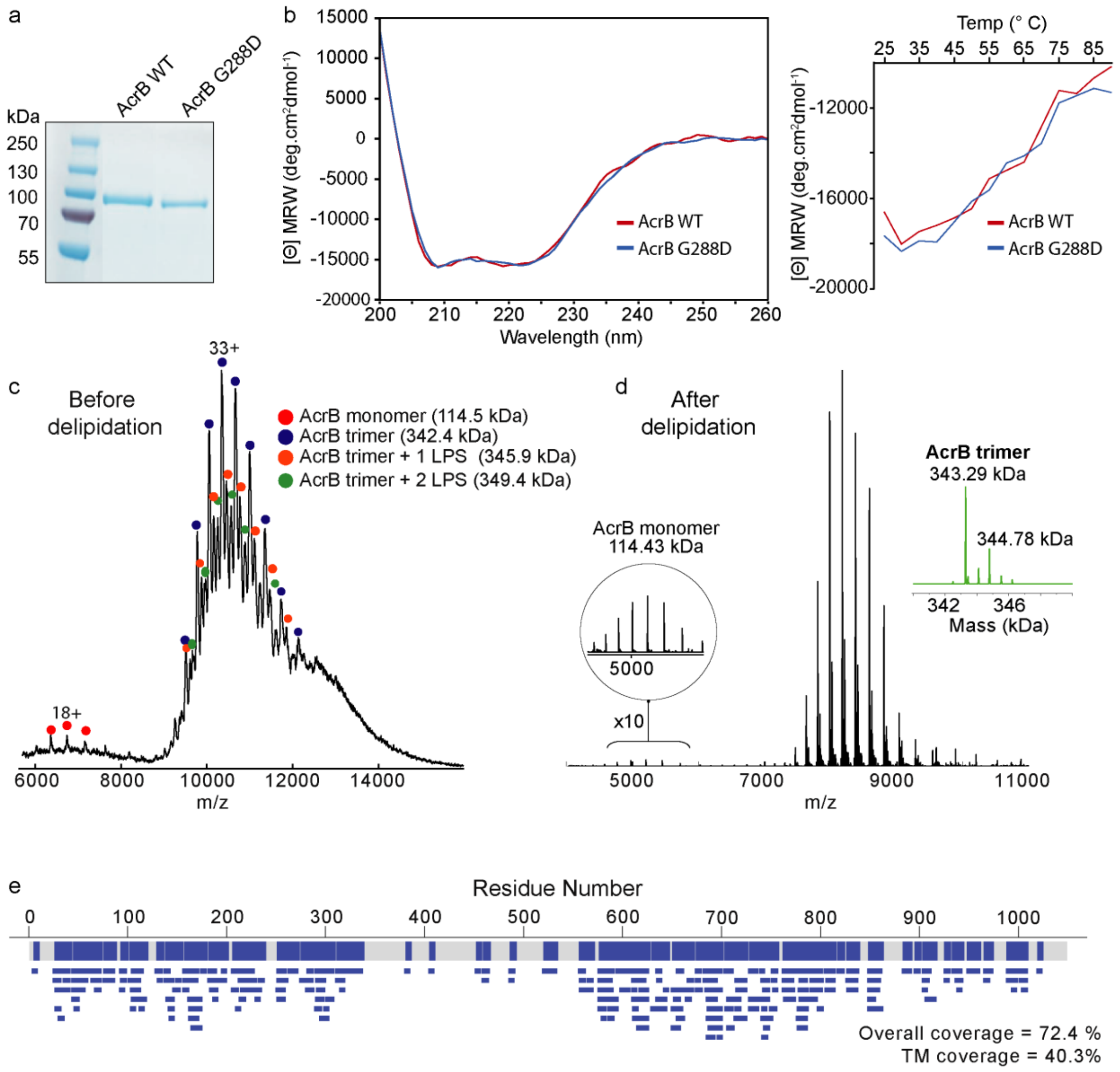


**Supplementary Figure 2. Biophysical characterization of AcrB$^{WT}$ and AcrB$^{G288D}$.** (**a**) SDS-PAGE of AcrB$^{WT}$ and AcrB$^{G288D}$ purified in DDM detergent micelles. (**b**) Circular dichroism and thermal melt (as determined by the loss of the helical signature at 222 nm) of AcrB reveals that the WT and G288D mutant have similar average secondary structure and thermal stability. Native mass spectra of AcrB before (**c**) and after (**d**) removal of detectable bound lipopolysaccharides (LPS) using optimized purification procedure as previously described (see Methods). (**e**) HDX-MS sequence coverage and peptide redundancy map of AcrB in DDM detergent micelles.

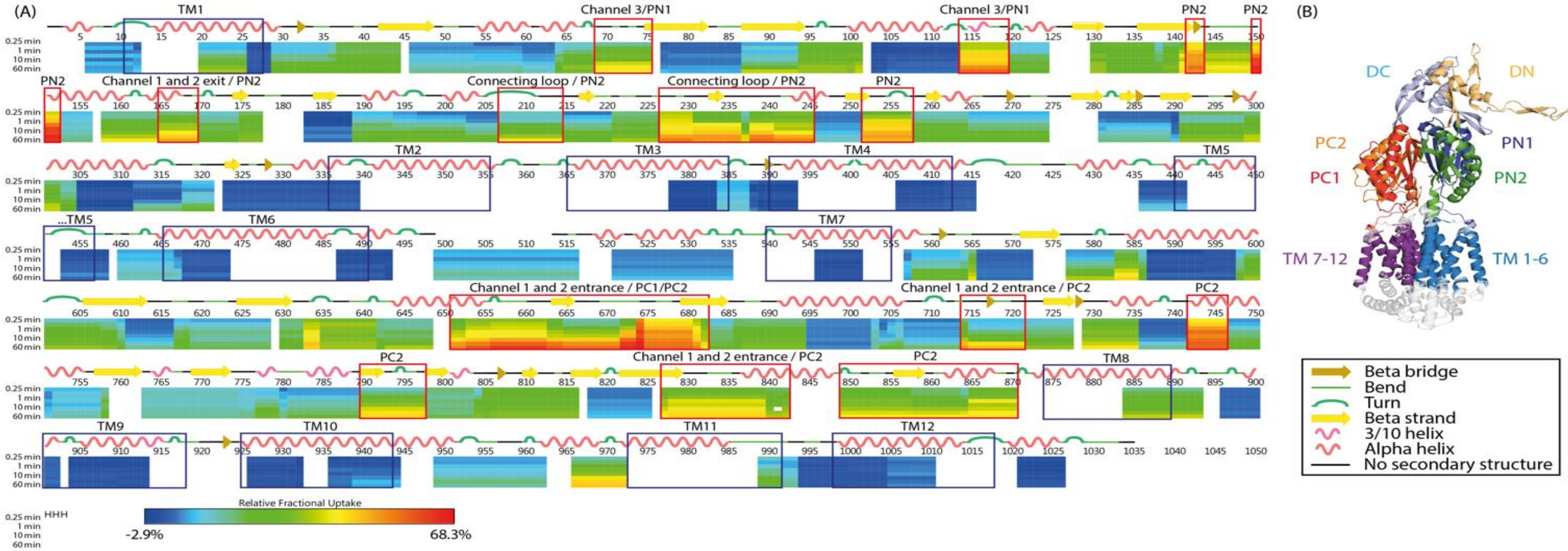


**Supplementary Figure 3. HDX-MS heat maps representing the relative fractional uptake of deuterium for peptides of AcrB.** The secondary structure of AcrB is shown above the peptide regions (key in box) and the subdomains of AcrB are shown on its monomer (right). The degree of relative fractional uptake of deuterium at each incubation time (from top to bottom: 15 s, 30s, 1 min, 3 min, 5 min, 10 min, 30 min and 60 min) are displayed according to the colour code shown. Uncoloured regions indicate areas with no peptide coverage. Red boxes annotate the regions that had the most significant change in uptake overtime. The secondary structure of AcrB is indicated above the heat map. All HDX-MS peptide data can be found in Supporting Data Table 1.

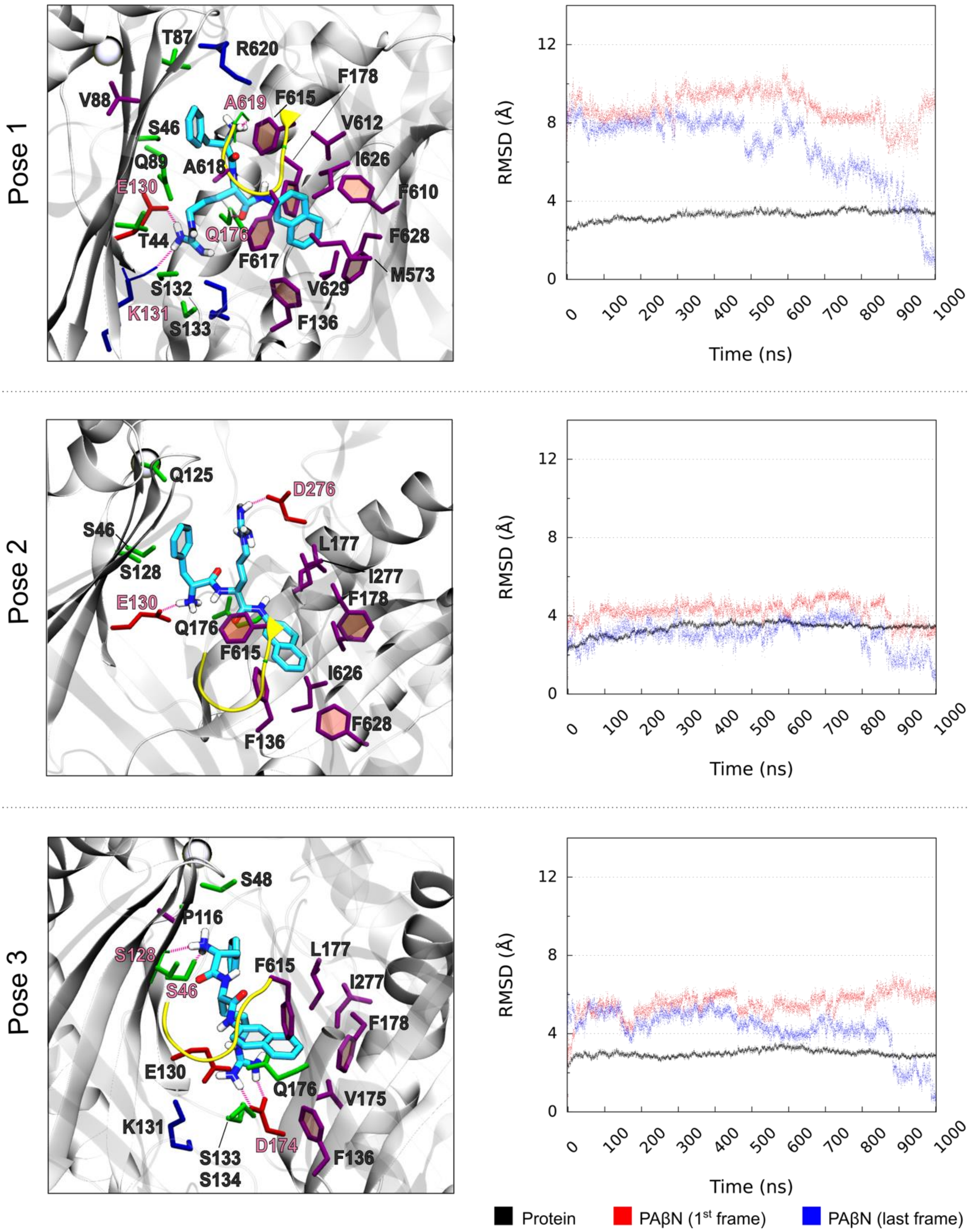


**Supplementary Figure 4. Representative binding poses and RMSDs of AcrB$^{WT}$-PAβN (Pose 1 discussed in the main text).** In the representation of the binding poses, PAβN is coloured by atom type (C atoms in cyan, N atoms in blue and oxygen atoms in red, H atoms in white – only polar H atoms are shown). Residues within 3.5 Å are also shown, coloured by residue type (red: acidic; blue: basic; green: polar; purple: hydrophobic). Hydrogen bonds formed by PAβN are highlighted through magenta sticks, and the involved residues are labelled in pink. The switch-loop is shown in yellow and the $C_\alpha$ atoms of the residues Q124 and Y758 belonging to the exit gate are represented as light blue spheres.

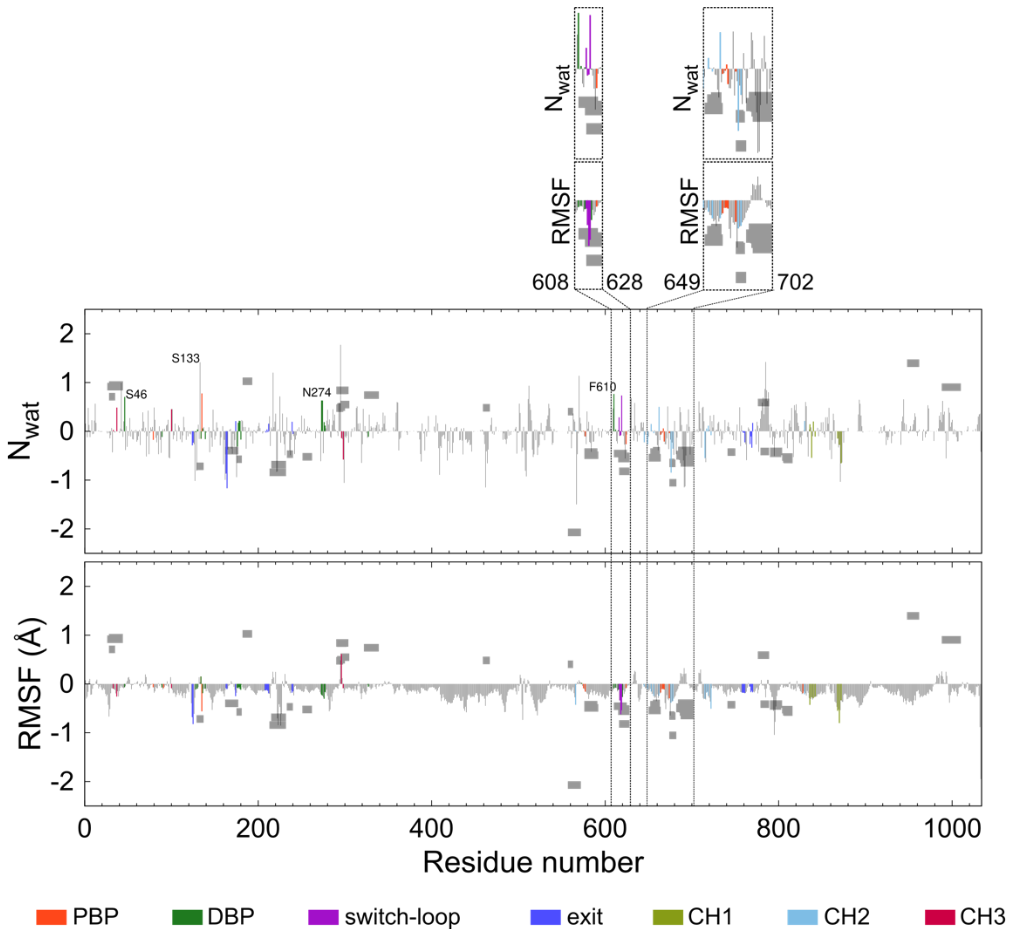


**Supplementary Figure 5. Difference in first hydration shell ($N_{wat}$) and RMSF between AcrB$^{WT}$-PAβN and apo AcrB$^{WT}$ (based on MD data from Pose 1 in Supplementary Fig. 4).** Differences in $N_{wat}$ and RMSF are represented as histograms, with regions directly involved in substrate transport highlighted in different colours (see Supplementary Table 1 for the definition of these regions). As a reference, HDX-MS data are represented as grey boxes (scale not shown). Both $N_{wat}$ and RMSF differences have been computed between the T monomer of AcrB$^{WT}$-PAβN and the L monomer of apo AcrB$^{WT}$ (see Methods). Regions of interest are highlighted in the upper part of the panel. In the $N_{wat}$ plot, labelled residues are directly involved in interactions with PAβN and have a higher hydration in AcrB$^{WT}$-PAβN than in apo AcrB$^{WT}$.

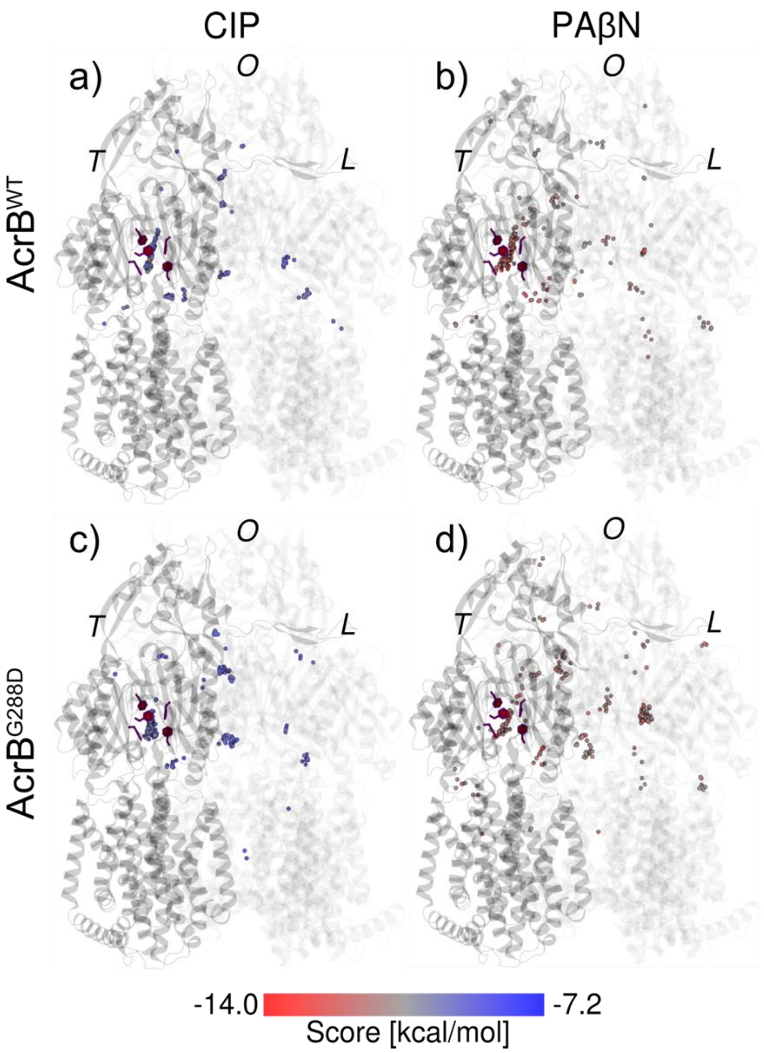


**Supplementary Figure 6. Distribution of top 200 docking poses (only the centres of mass are shown for clarity) for CIP and PAβN onto AcrB$^{WT}$ (a,b) and AcrB$^{G288D}$ (c, d).** The spheres are coloured according to the value of the (pseudo) free-energy of binding (Score). The monomers L, T and O are shown as transparent ribbons (T and O darkest and lightest, respectively). These distributions and the data in Supplementary Tables 2-3 justify our focus on the DBP of the T monomer. Details of the docking protocol are given in Methods.

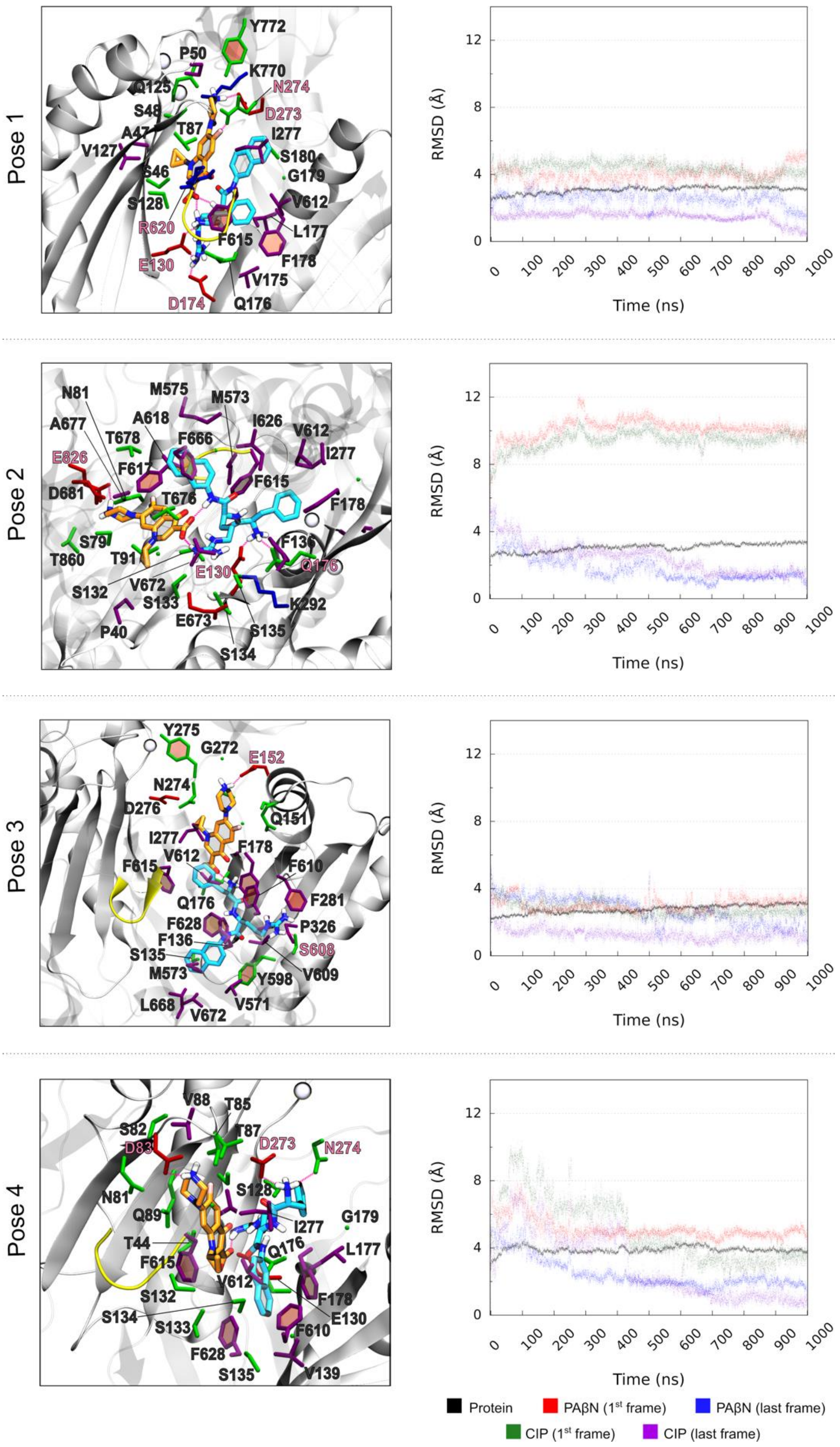


**Supplementary Figure 7. Representative binding poses and RMSDs of AcrB$^{WT}$-CIP-PAβN (Pose 1 discussed in the main text).** To distinguish between the inhibitor and antibiotic, carbon atoms of CIP and PAβN are coloured in orange and cyan, respectively. See Supplementary Fig. 4 for further details.

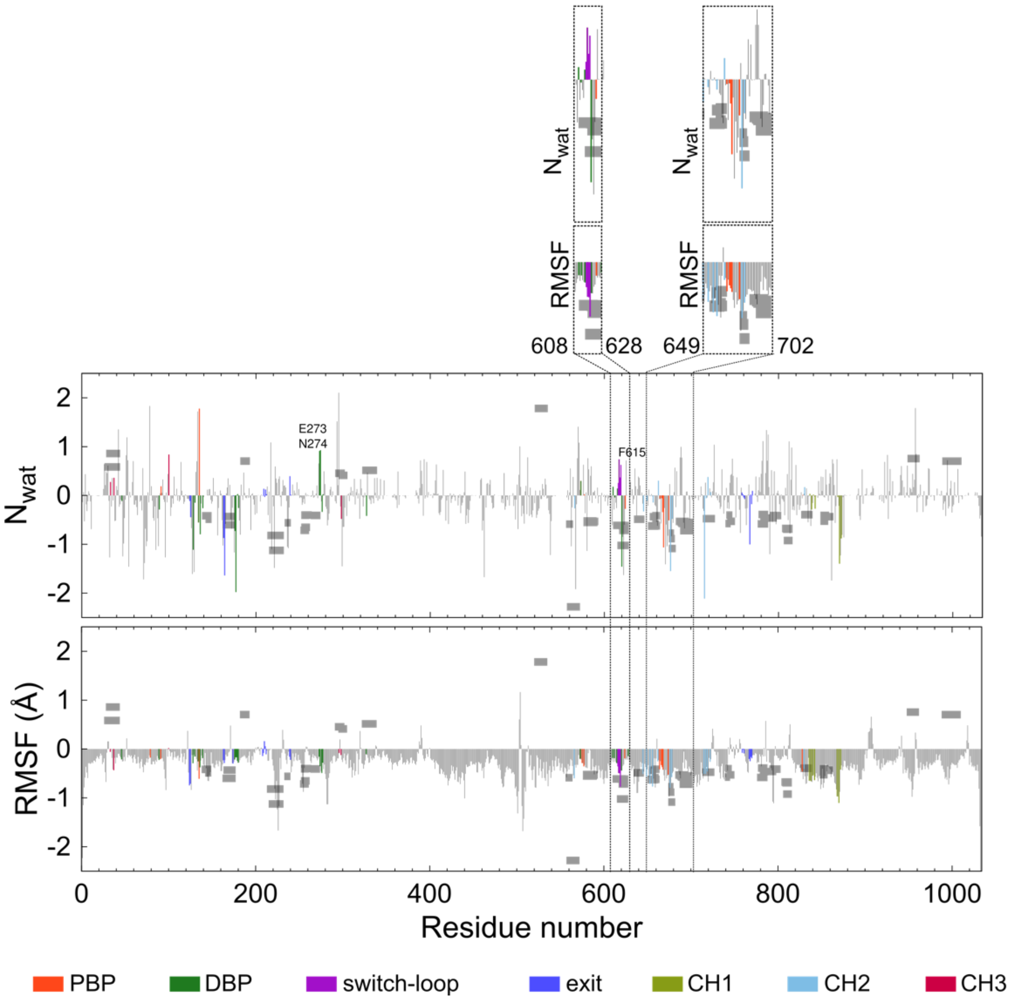


**Supplementary Figure 8. Difference in first hydration shell ($N_{wat}$) and RMSF between AcrB$^{WT}$-CIP-PAβN and apo AcrB$^{WT}$ (based on MD data from Pose 1 in Supplementary Fig. 4).** See Supplementary Fig. 5 for details.

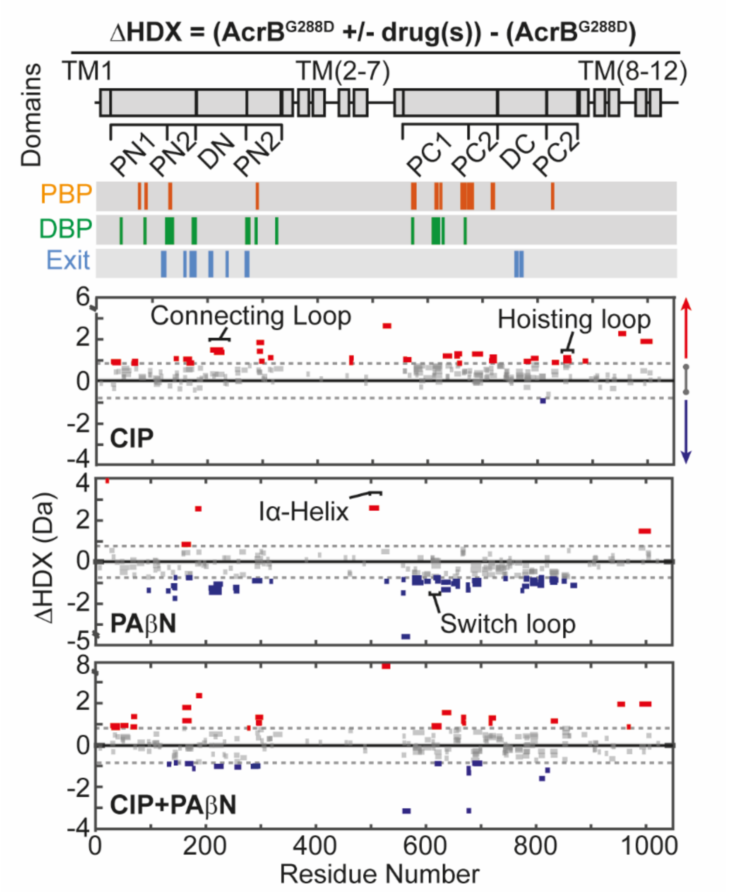


**Supplementary Figure 9. Influence of drugs on AcrB$^{G288D}$ structural dynamics**. Sum differential HDX (ΔHDX) plots for different drug conditions (ΔHDX = (AcrB$^{G288D}$+drug(s)) – AcrB$^{G288D}$) for all time points collected. All data reported as in Fig. 2 and 4. All HDX-MS peptide data can be found in Supporting Data Table 2.

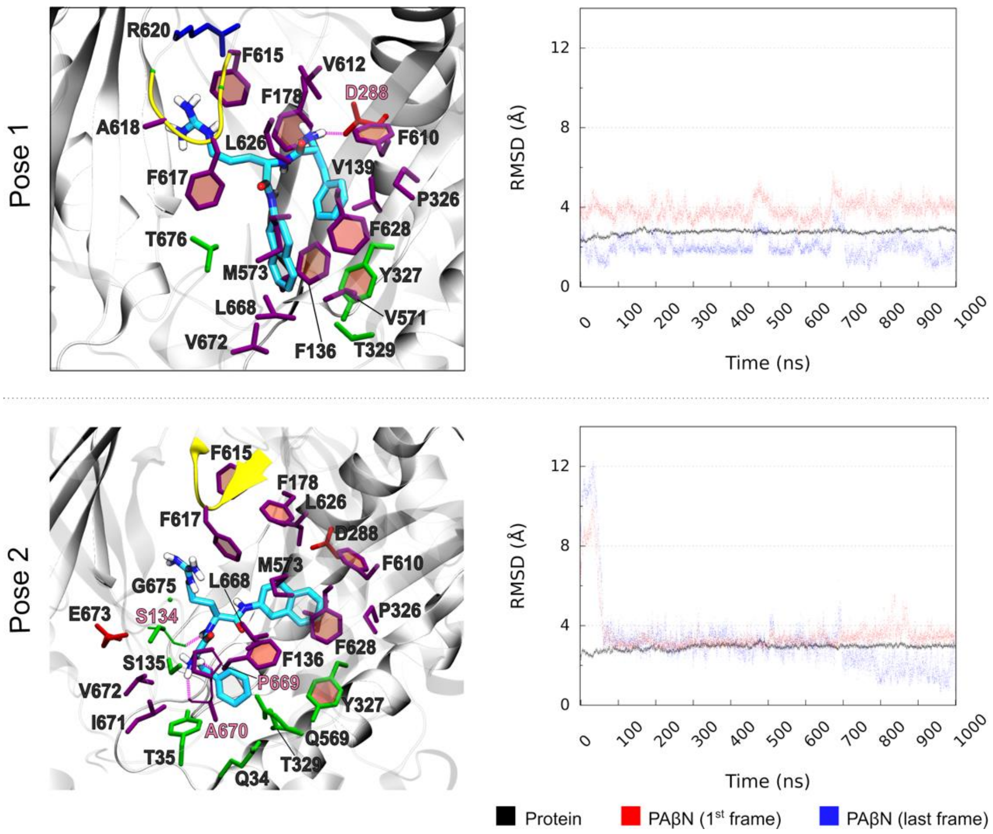


**Supplementary Figure 10. Representative binding poses and RMSDs of AcrB$^{G288D}$-PAβN (Pose 1 discussed in the main text).** See Supplementary Fig. 4 for details.

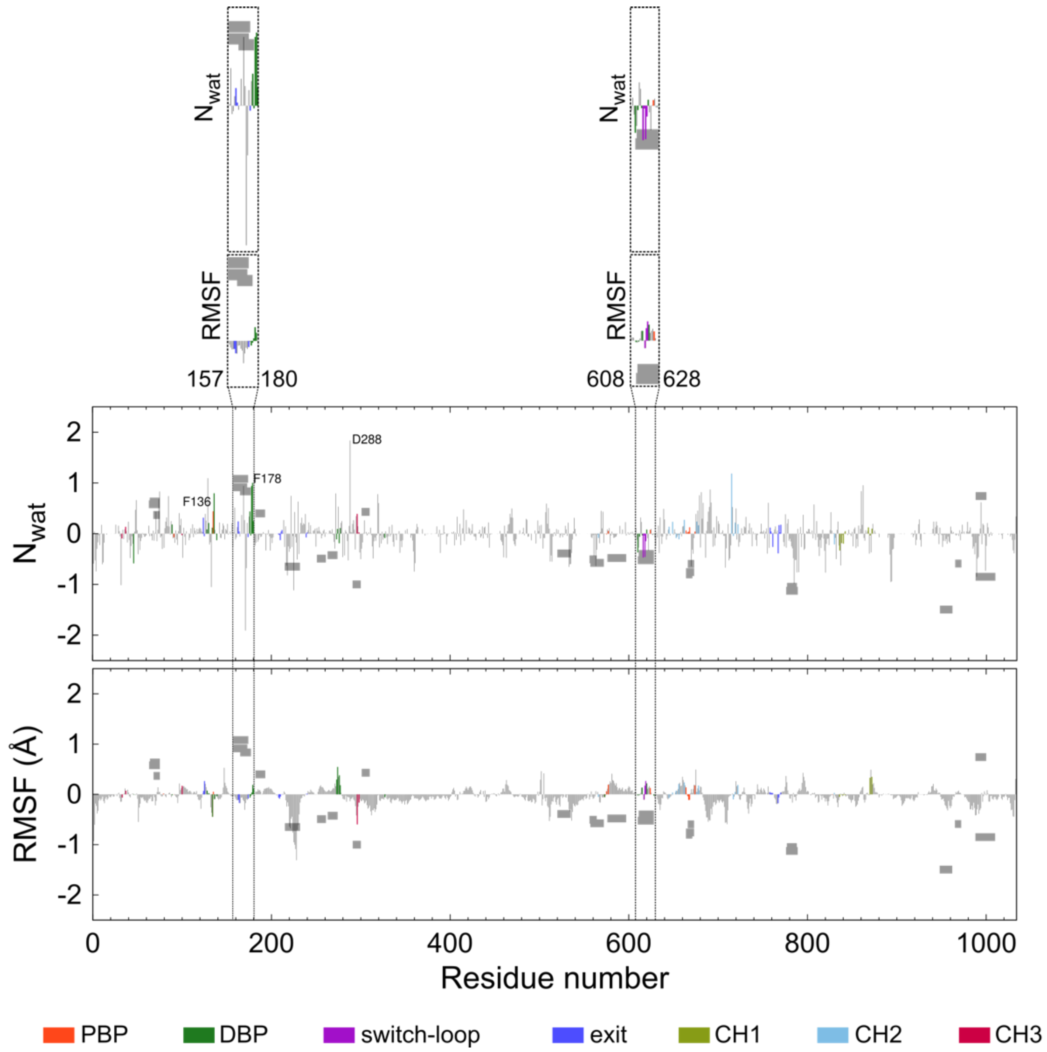


**Supplementary Figure 11.** Difference in first hydration shell ($N_{wat}$) and RMSF between AcrB$^{G288D}$-PAβN and apo AcrB$^{WT}$-PAβN (based on MD data from Pose 1 in Supplementary Fig. 10). Both AcrB$^{G288D}$-PAβN and AcrB$^{WT}$-PAβN were considered in the T state (see Methods). See Supplementary Fig. 5 for details.

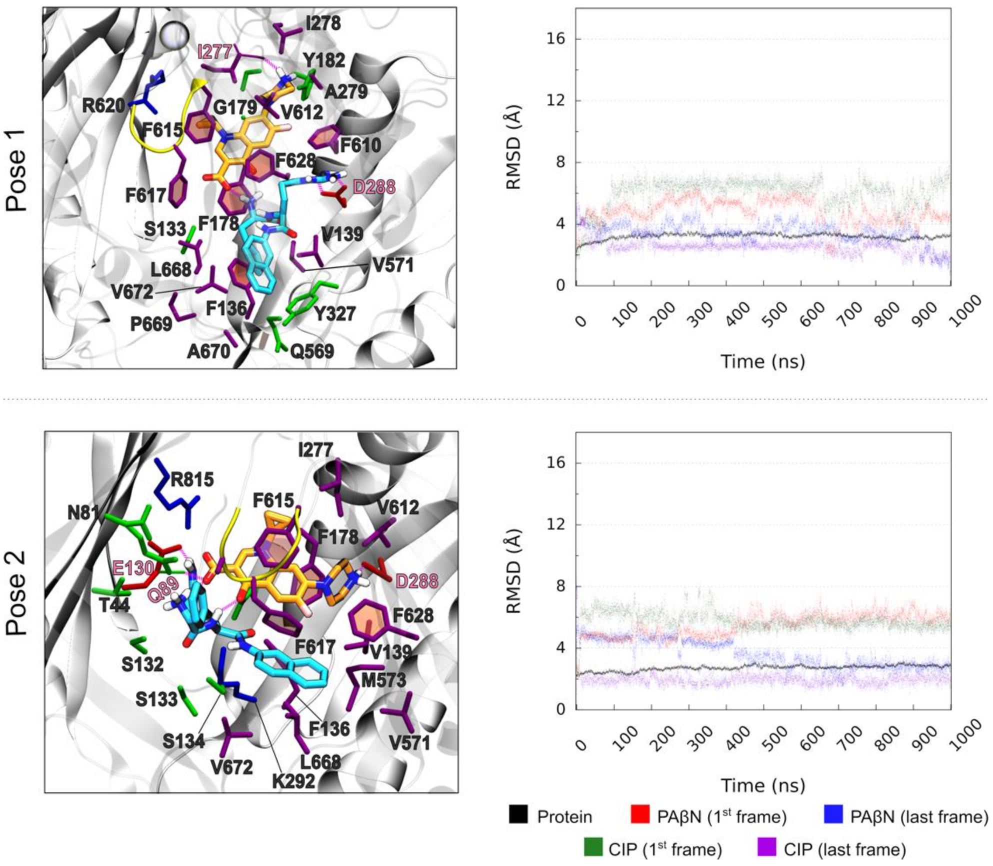


**Supplementary Figure 12. Representative binding poses and RMSDs of AcrB$^{G288D}$-CIP-PAβN (Pose 1 discussed in the main text).** See Supplementary Fig. 4 for details.

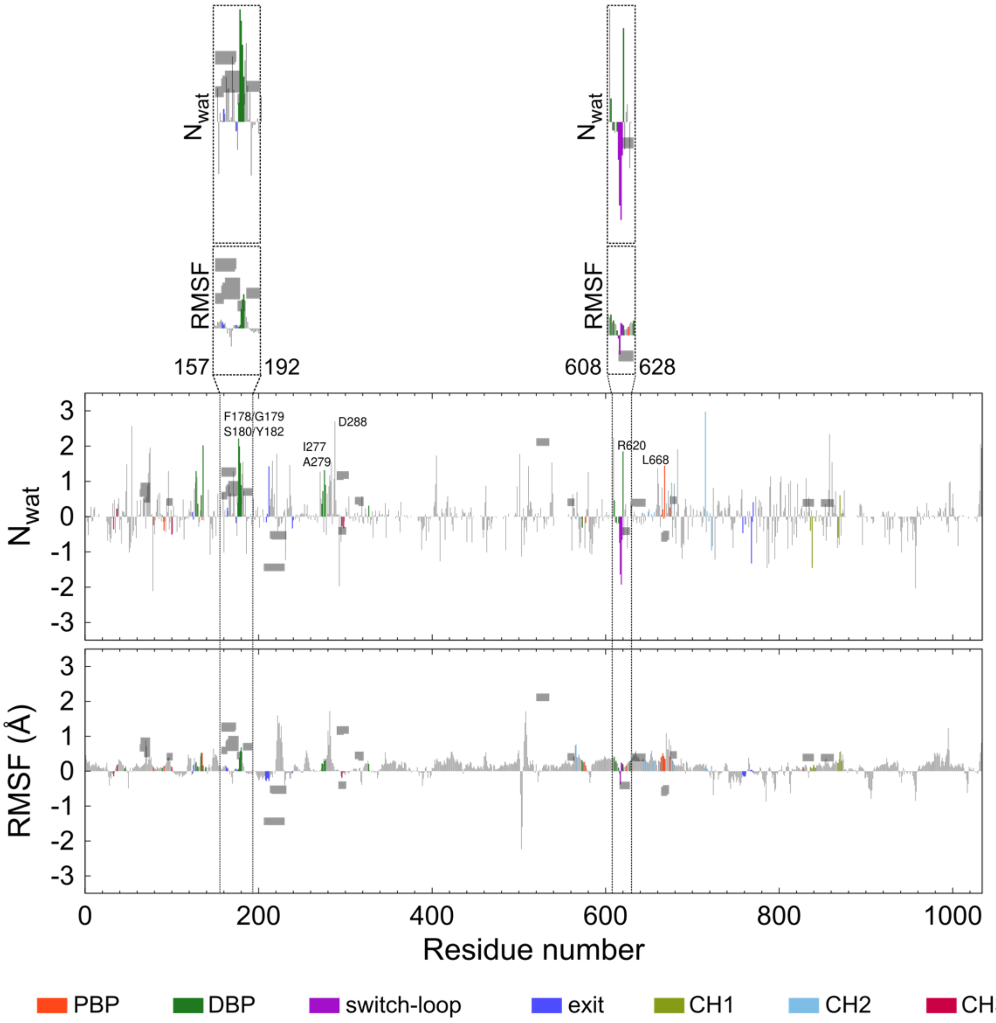


**Supplementary Figure 13. Difference in first hydration shell ($N_{wat}$) and RMSF between AcrB$^{G288D}$-CIP-PAβN and apo AcrB$^{WT}$-CIP-PAβN (based on MD data from Pose 1 in Supplementary Fig. 12).** Both AcrB$^{G288D}$-CIP-PAβN and AcrB$^{WT}$-CIP-PAβN were considered in the T state (see Methods). See Supplementary Fig. 5 for details.

**Supplementary Table 1.** List of residues considered in the regions of AcrB.

| Regions | Residues |
|---|---|
| Central cavity | 25-33, 36, 37, 96, 97, 385-389, 457-466, 468, 469 |
| Distal binding pocket (DBP) | 44, 46, 48, 87, 89, 128, 130, 132, 134, 136, 139, 176--180, 273, 274, 276, 277, 327, 573, 610, 612, 615, 617, 620, 626, 628 |
| Hydrophobic trap (HT) | 136, 178, 289, 291, 573, 610, 612, 615, 626, 628 |
| Proximal binding pocket (PBP) | 79-81, 89-91, 132-134, 573, 575, 577, 617, 662-669, 672-681, 683, 717, 719, 815, 826, 828-830 |
| Connecting-loop | 206-243 |
| Iα-Helix | 520-534 |
| Switch-loop | 615-620 |
| Hosting-loop | 860-871 |
| Exit gate (EG) | 124, 125, 163, 164, 174, 208-221, 239, 240, 758-761, 767-770 |
| Channel 1 (CH1) | 836, 838, 840, 842, 868, 870, 872 |
| Channel 2 (CH2) | 566, 645, 649, 653, 656, 662, 676, 678, 715, 717, 719, 722, 830 |
| Channel 3 (CH3) | 33, 37, 100, 296, 298 |

| | %res | 30% | | | 40% | | |
|---|---|---|---|---|---|---|---|
| | **Site** | **N** | **$\Delta G_{max}$** | **<$\Delta G$>** | **N** | **$\Delta G_{max}$** | **<$\Delta G$>** |
| **PAβ N** | **$PBP_L$** | 12 | -11.5 | -10.9+-0.3 | 3 | -11.3 | -10.9+-0.3 |
| | **$PBP_T$** | 19 | -13.0 | -11.6+-0.6 | - | - | - |
| | **$CH1_L$** | 1 | -10.8 | -10.8 | 1 | -10.8 | -10.8 |
| | **$CH1_T$** | - | - | - | - | - | - |
| | **$CH2_L$** | 1 | -11.3 | -11.3 | - | - | - |
| | **$CH2_T$** | - | - | - | - | - | - |
| | **$CH3_L$** | 15 | -12.3 | -11.3+-0.4 | 9 | -12.3 | -11.4+-0.4 |
| | **$CH3_T$** | 33 | -12.5 | -11.4+-0.4 | 22 | -12.5 | -11.5+-0.4 |
| | **$DBP_T$** | 148 | -13.7 | -11.6+-0.7 | 87 | -13.7 | -11.7+-0.7 |
| **CIP** | **$PBP_L$** | - | - | - | - | - | - |
| | **$PBP_T$** | - | - | - | - | - | - |
| | **$CH1_L$** | - | - | - | - | - | - |
| | **$CH1_T$** | - | - | - | - | - | - |
| | **$CH2_L$** | - | - | - | - | - | - |
| | **$CH2_T$** | - | - | - | - | - | - |
| | **$CH3_L$** | 55 | -10.2 | -9.4+-0.3 | 19 | -9.6 | -9.3+-0.2 |
| | **$CH3_T$** | 31 | -10.0 | -9.5+-0.2 | 4 | -9.7 | -9.4+-0.3 |
| | **$DBP_T$** | 123 | -11.5 | -9.7+-0.4 | 20 | -10.3 | -9.6+-0.4 |

**Supplementary Table 2.** Number of poses, maximum and average (pseudo)free-energy of binding in $PBP_L$, $PBP_T$, $CH1_L$, $CH1_T$, $CH2_L$, $CH2_T$, $CH3_L$, $CH3_T$ and $DBP_T$ of $AcrB_{WT}$ as obtained from blind ensemble docking (for each compound we docked 10 conformations onto 10 conformations of the protein, see Methods for details). The percentages in the first row are meant to identify the poses having contacts (that is minimum ligand-residue distance below a cutoff – here set to 3.5 Å) at least with 30% or 40% of residues lining the corresponding site. From blind docking calculations it is clear that, overall, there is a significant overlap between the poses of PAβN and CIP (see Supplementary Fig. 6a,b). Both compounds accumulate high-affinity binding poses within the DBP of monomer T ($DBP_T$). It is also evident how the inhibitor has a larger binding affinity than CIP towards that pocket. However, other poses for PAβN are found at $CH1_{L/T}$, $CH2_L$, and $CH3_{L/T}$, as well as within $PBP_{L/T}$. Regarding CIP, we also see poses just beneath the CH3 of monomers L and T and farther away from CH1 of monomer T, while as expected no poses are found within the PBP. These data suggest that PAβN, whose affinities at $CH3_{L/T}$ are comparable to those at $DBP_T$, could in principle compete with CIP also during the uptake by AcrB, thus not only regarding binding at the preferred site.

| | %res | 30% | | | 40% | | |
|---|---|---|---|---|---|---|---|
| | Site | N | ΔG$_{max}$ | <ΔG> | N | ΔG$_{max}$ | <ΔG> |
| PAβ N | PBP$_L$ | - | - | - | - | - | - |
| | PBP$_T$ | 3 | -11.5 | -11.1+-0.3 | - | - | - |
| | CH1$_L$ | - | - | - | - | - | - |
| | CH1$_T$ | 2 | -10.8 | -10.8+-0.0 | 2 | -10.8 | -10.8+-0.0 |
| | CH2$_L$ | - | - | - | - | - | - |
| | CH2$_T$ | 2 | -11.1 | -11.1+-0.0 | - | - | - |
| | CH3$_L$ | 18 | -12.9 | -11.5+-0.6 | 7 | -12.0 | -11.5+-0.4 |
| | CH3$_T$ | 11 | -13.5 | -11.7+-0.8 | 4 | -11.8 | -11.3+-0.3 |
| | DBP$_T$ | 24 | -13.4 | -11.6+-0.7 | 2 | -11.3 | -11.2+-0.1 |
| CIP | PBP$_L$ | - | - | - | - | - | - |
| | PBP$_T$ | - | - | - | - | - | - |
| | CH1$_L$ | - | - | - | - | - | - |
| | CH1$_T$ | - | - | - | - | - | - |
| | CH2$_L$ | - | - | - | - | - | - |
| | CH2$_T$ | - | - | - | - | - | - |
| | CH3$_L$ | 14 | -9.9 | -9.2+-0.3 | - | - | - |
| | CH3$_T$ | 9 | -9.4 | -9.1+-0.2 | - | - | - |
| | DBP$_T$ | 5 | -9.8 | -9.5+-0.2 | 1 | -9.3 | -9.3+-0.0 |

**Supplementary Table 3.** Number of poses, maximum and average (pseudo)free-energy of binding in PBP$_L$, PBP$_T$, CH1$_L$, CH1$_T$, CH2$_L$, CH2$_T$, CH3$_L$, CH3$_T$ and DBP$_T$ of AcrB$^{G288D}$ as obtained from blind ensemble docking (for each compound we docked 10 conformations onto 10 conformations of the protein, see caption of Supplementary Table 2 for further details and Supporting Fig. 6c,d for the distribution of the poses).

| System | Reference |
|---|---|
| AcrB$^{WT}$-PAβN (T state) | Apo AcrB$^{WT}$ (L state) |
| AcrB$^{WT}$-CIP-PAβN (T state) | Apo AcrB$^{WT}$ (L state) |
| AcrB$^{G288D}$-PAβN (T state) | AcrB$^{WT}$-PAβN (T state) |
| AcrB$^{G288D}$-CIP-PAβN (T state) | AcrB$^{G288D}$-CIP-PAβN (T state) |

**Supplementary Table 4.** Systems considered for the analyses of flexibility and hydration properties (based on MD simulations), and respective reference structures.
